# Microtubule plus-end dynamics are tightly coupled to the turnover of the MOR1 polymerase

**DOI:** 10.1101/2024.10.30.621176

**Authors:** Ryan C Eng, Laryssa S Halat, Chak-Chung Kuo, Aida Rakei, Cecily D Costain, Kamryn A Diehl, Ron Blutrich, Bettina Lechner, Yi Zhang, Ankit Walia, Jose M Alonso, Daniel Coombs, Geoffrey O Wasteneys

**Affiliations:** Department of Botany, University of British Columbia, Vancouver, BC V6T 1Z4, Canada; Department of Plant and Microbial Biology, Program in Genetics, North Carolina State University, Raleigh, NC 27695, USA; Department of Mathematics, University of British Columbia, Vancouver, BC V6T 1Z2, Canada

**Author notes:** These authors (R.C.E., L.S.H., C.C.K.) contributed equally to the manuscript. Corresponding author: Geoffrey Wasteneys.

## Abstract

MOR1/XMAP215/Dis1/Stu2 family proteins function as polymerases that promote microtubule assembly through tubulin-interacting N-terminal tumor-overexpressed gene (TOG) domains. It remains unclear how the polymerase activity of these proteins is coordinated with their movement along the growing microtubule lattice. Previously, it was found that a L174F mutation in the N-terminal TOG domain of the 217 kDa *Arabidopsis thaliana* MICROTUBULE ORGANIZATION 1 (MOR1) protein impaired microtubule growth and shrinkage rates in a temperature-dependent manner. In this study, we used a homologous recombination strategy to generate stable transgenic lines expressing full-length MOR1 and mutant mor1-1^L174F^ proteins tagged with the yellow fluorescent protein YPet. When expressed in *mor1* mutant backgrounds, MOR1-3xYPet showed strong association with both the growing and shrinking microtubule plus ends, consistent with its involvement in both polymerization and depolymerization. By contrast, when expressed in a wild-type background, MOR1-3xYPet was redistributed along the microtubule lattice, microtubule growth rates were reduced, and plant growth was stunted. By quantifying fluorescence recovery after photobleaching, we confirmed that MOR1 has greater affinity for the plus end than the lattice. Exploiting the FRAP technique further, we identified a positive and linear correlation between microtubule polymerization rates and MOR1 turnover at the microtubule plus end. When microtubule growth rates were reduced in the mor1-1-YPet line by shifting the temperature from 21 to 30°C and for MOR1-3xYPet under taxol treatments, MOR1’s turnover was delayed. In contrast, microtubule growth rates and MOR1 turnover were hastened at 30°C or after CRISPR/Cas9-mediated knock out of the CLASP protein. These findings support a new model whereby MOR1 has finite processivity on the microtubule, catalyzing a limited fixed number of tubulin additions before dissociation.

**HIGHLIGHTS:** MOR1 has high affinity for both growing and shrinking microtubule plus ends, consistent with a function in both polymerization and depolymerization events.

MOR1 has greater affinity for microtubule plus ends than the lattice, and lattice binding associated with overexpression is detrimental to plant growth.

MOR1 transiently associates with the microtubule plus end, the duration of which is tightly coupled with its polymerase activity.

## INTRODUCTION

Microtubules, the dynamic filamentous polymers that drive cell division, motility, and intracellular transport in eukaryotic cells, are under the tight control of microtubule-associated proteins (MAPs). MICROTUBULE ORGANIZATION 1 (MOR1) is a MAP first identified from a forward genetics screen in the model plant *Arabidopsis thaliana* (Whittington et al., 2001). Its indispensable role in microtubule dynamics during plant growth and development is evident from the drastic growth and microtubule defects of the temperature-sensitive *mor1-1* mutant allele (Whittington et al., 2001) and the embryo-lethal phenotypes of the *gem1-1* mutant allele of *MOR1* (Twell et al., 2002). Understanding the mechanism by which MOR1 facilitates microtubule assembly is at the essence of understanding microtubule dynamics and essential plant developmental processes.

MOR1 is a member of the XMAP215/Dis1 family proteins characterized by their N-terminal TOG (tumour <u>o</u>verexpressed <u>g</u>ene) domains, which generally contain six HEAT (<u>H</u>untingtin, <u>E</u>longation factor 3, protein phosphatase 2<u>A</u>, and <u>T</u>OR1) repeats folded into side-by-side α-helices connected by intra-HEAT structures (Al-Bassam et al., 2007; Al-Bassam & Chang, 2011; Lechner et al., 2012; Slep & Vale, 2007). These TOG domain proteins function as microtubule polymerases (Brouhard et al., 2008; Podolski et al., 2014; Van Breugel et al., 2003) but also promote microtubule depolymerization under some conditions (Shirasu-Hiza et al., 2003; Vasquez et al., 1994, 1999). MOR1’s polymerase function is supported by the observations that at the restrictive temperature, the *mor1-1* mutant has short microtubules (Kawamura et al., 2006; Whittington et al., 2001) that grow at reduced rates (Kawamura & Wasteneys, 2008). Interestingly, microtubule shrinkage rates are also significantly reduced in *mor1-1* mutants (Kawamura & Wasteneys, 2008), suggesting that MOR1 also supports rapid microtubule depolymerization.

Consistent with their polymerase activity, several XMAP215/Dis1 proteins including XMAP215 (Brouhard et al., 2008), Stu2 (Van Breugel et al., 2003), and Alp14 (Al-Bassam et al., 2012; Garcia et al., 2001) are associated with the plus ends of microtubules. However, MOR1 localization studies have remained inconclusive. One immunolabelling study indicated increased labelling at the midzone of the mitotic spindles and phragmoplasts but the antibody also showed a weak but punctate distribution along interphase microtubules (Twell et al., 2002). A separate study using a different antibody identified MOR1 epitopes along the entire lengths of interphase, spindle and phragmoplast microtubules (Kawamura et al., 2006). According to live-cell imaging, a fluorescent reporter of the first two TOG domains of MOR1 (TOG1-TOG2), which was capable of polymerizing microtubules *in vitro* and rescuing the *mor1-1* mutant phenotype, similarly distributed along the full length of microtubules (Lechner et al., 2012). These results put MOR1 at odds with its XMAP215/Dis1 homologues that showed microtubule plus-end tracking.

To catalyze microtubule polymerization, the N-terminal TOG domains of XMAP215/Dis1 proteins need to distinguish between free and polymerized tubulin (Ayaz et al., 2012). Evidence suggests that this is achieved through diverse binding specificity of TOG domains. For instance, the first two TOG domains of XMAP215 can bind free tubulin (Widlund et al., 2011) and the fifth TOG domain of *Drosophila melanogaster* Msps can bind the microtubule lattice (Byrnes & Slep, 2017). Unlike XMAP215, the first two TOG domains of MOR1 showed relatively low affinity for free tubulin and relatively high affinity for the microtubule lattice *in vitro* (Lechner et al., 2012), suggesting that MOR1’s mode of function differs from that of its animal and fungal homologues. A conserved microtubule-binding motif at the N-terminus of MOR1, which is found closer to the C-terminus in XMAP215 and Msps, could explain this difference in affinity for the microtubule lattice (Currie et al., 2011; Lechner et al., 2012; Widlund et al., 2011).

Studies of the *mor1-1* temperature-sensitive mutation highlight the importance of MOR1’s first TOG domain in microtubule polymerization (Kawamura et al., 2006; Kawamura & Wasteneys, 2008; Whittington et al., 2001). Immunolabelling experiments demonstrated that the mor1-1 protein (with the L174F mutation) remains associated with microtubules at the restrictive temperature (> 28°C), indicating that its inhibitory effects are not the result of weakened association with microtubules (Kawamura et al., 2006). Indeed, subsequent microtubule co-sedimentation assays indicated that the L174F substitution in a HEAT repeat increases MOR1’s affinity for the microtubule lattice *in vitro* (Lechner et al., 2012).

Recognizing how an amino acid substitution in TOG1 can provide valuable insight into the relationship between polymerase activity and processivity, we produced stable transgenic lines expressing full-length wild-type MOR1 and mor1-1 mutant proteins tagged with a yellow fluorescent protein. To overcome the challenge of cloning and modifying the *MOR1* gene, we used recombineering (Zhou et al., 2011) to insert the fluorescent protein-encoding sequences and create the *mor1-1* point mutation in the *MOR1* gene. We confirmed that MOR1 associates with microtubule arrays at all stages of the cell cycle and development, and found that it strongly associates with both the growing and shrinking plus ends of microtubules. We also developed a Fluorescence Recovery After Photobleaching (FRAP) technique to measure MOR1 turnover at the plus end and the lattice, to compare the turnover of wild-type MOR1 and the polymerase-defective mor1-1 protein, and to quantify how perturbations to microtubule dynamics affect MOR1’s turnover at the plus end.

## RESULTS

### Recombineering MOR1 and mor1-1 fluorescent reporters

Our previous attempts to establish a system to study MOR1 *in vivo* were met with several major challenges including *MOR1’s* large size (14 kb) and complex gene structure, with 53 exons (Figure 1A). Recombineering exploits homologous recombination in transformable bacterial artificial chromosomes to enable reporter construct design for genes like *MOR1* (Zhou et al., 2011). Using this technique, we generated endogenous promoter-driven translational reporters of the wild-type *MOR1* and the temperature-sensitive allele *mor1-1*, a single nucleotide mutation (Figure 1A) leading to a L174F substitution (Figure 1B). The wild-type MOR1 was tagged at the C-terminus with 3 copies of the yellow fluorescent protein YPet (*MOR1_pro_:MOR1-3xYPet*, Figure 1B), and mor1-1 was tagged with a single YPet (*MOR1_pro_:mor1-1-YPet*, Figure 1B).

**Figure 1.**
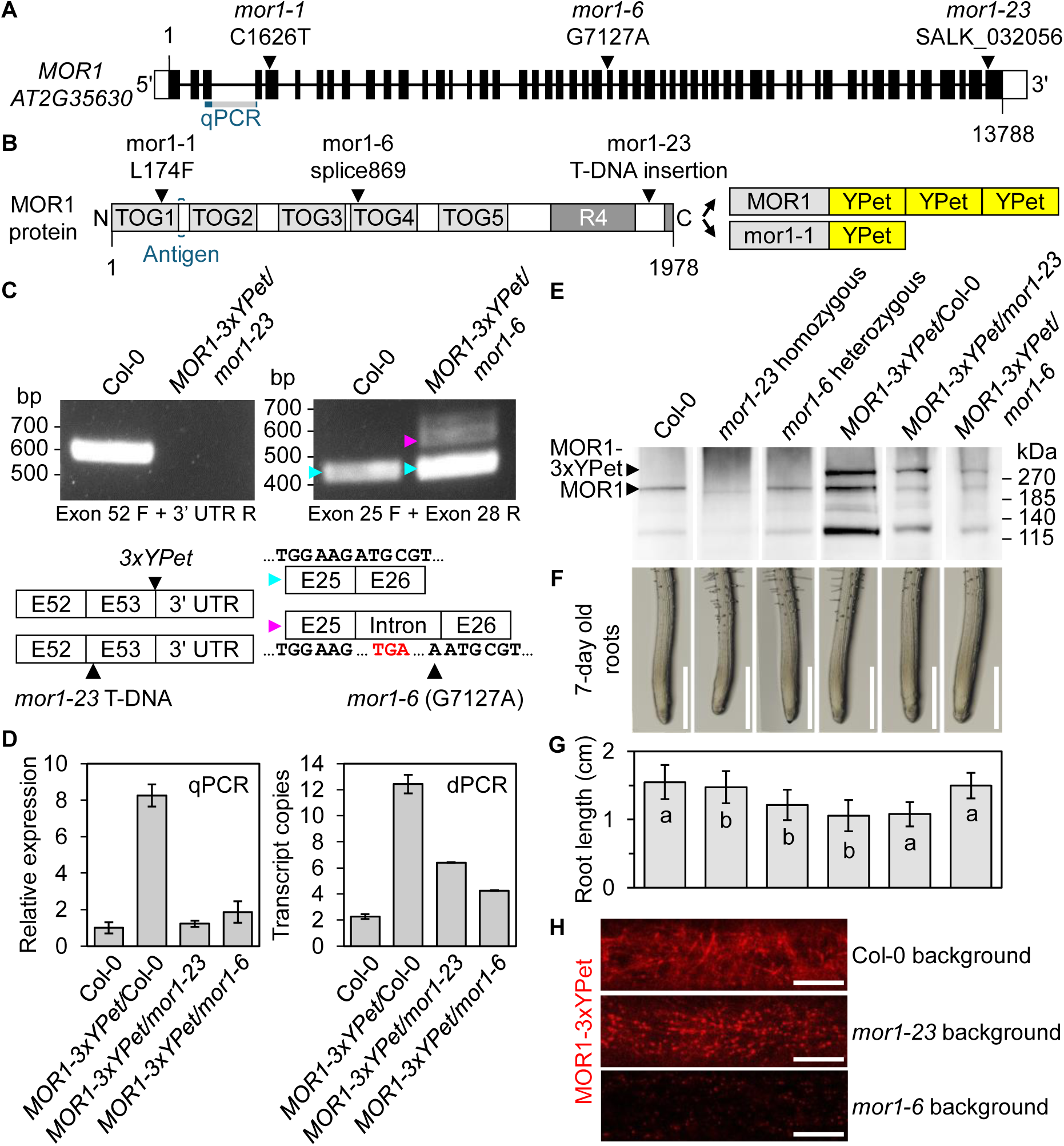
Molecular and phenotypic analyses of the *MOR1-3xYPet* constructs in the wild-type (Col-0) background and in the *mor1-6* and *mor1-23* mutant backgrounds. **(A)** Structure of the *Arabidopsis MOR1* genomic sequence. Three *MOR1* mutant alleles are shown: *mor1-1* (C1626T), *mor1-6* (G7127A), and *mor1-23* (SALK_032056). Black boxes represent exons, horizontal lines represent introns, and white boxes represent 5’ and 3’ untranslated regions (UTRs). qPCR primers were designed to amplify a region spanning exons 3 and 4 (marked in blue). **(B)** The MOR1 protein contains five TOG domains and a conserved C-terminal R4 domain. Mutant proteins include mor1-1^L174F^, mor1-6 (splice mutation before amino acid 869), and mor1- 23 (T-DNA insertion). The linker between TOG1 and TOG2 was used as the antigen for creating the MOR1 antibody (Kawamura et al., 2006). Two C-terminal fusion proteins were generated: MOR1-3xYPet (three copies of YPet attached to MOR1), and mor1-1-YPet (one copy of YPet attached to mor1-1^L174F^). **(C)** RT-PCR analysis of *mor1-23* and *mor1-6* mutant backgrounds. For *mor1-23*, primers failed to amplify through the *3xYPet* sequence or the *mor1-23* T-DNA insertion. For *mor1-6*, blue arrows indicate wild-type cDNA amplicons when introns are correctly spliced, and magenta arrows indicate amplicons longer than wild-type cDNA but shorter than genomic DNA due to a retained intron. Forward and reverse primer binding sites are indicated below each gel. Exon-intron structures are confirmed by sequencing and marked with arrows corresponding to the amplicons. The premature stop codon within the intron is marked red. **(D)** RT-qPCR and digital PCR (dPCR) analysis of *MOR1* transcript levels when *MOR1-3xYPet* was expressed in Col-0, *mor1-23*, and *mor1-6* backgrounds. Primers bind exons 3 and 4 as shown in **(A)**. Bars and error bars represent mean values of 3 technical replicates ± SE. **(E)** Western blot analysis of MOR1 proteins in seedling protein extracts. The affinity-purified MOR1 polyclonal antibody labelled 218 kDa bands (endogenous MOR1 protein), 300 kDa bands (MOR1-3xYPet), and bands with lower molecular weights (approx. 125 and 160 kDa). **(F)** Representative images showing root tip morphologies of 7-day old seedlings including wild-type Col-0, homozygous *mor1-23*, heterozygous *mor1-6*, and *MOR1-3xYPet* expressed in Col-0, *mor1-23*, and *mor1-6* backgrounds. Scale bar represents 0.5 mm. Sample sizes: 14-16 seedlings. **(G)** Mean root length (in cm) of genotypes shown in **F**. Bars and error bars represent mean ± SD. Kruskal-Wallis *p* = 3.69E-9. Letters represent statistical differences between genotypes revealed by Dunn’s test. **(H)** Confocal micrographs of MOR1-3xYPet in wild-type, *mor1-23*, and *mor1-6* backgrounds, as seen in etiolated hypocotyl cells. MOR1-3xYPet is more tightly localized to microtubule tips in the *mor1-23* and *mor1-6* backgrounds while distributed along the microtubule lattice in the wild-type background. Scale bars represent 10 μm.

Since *MOR1* is an essential single-copy gene (Whittington et al., 2001), total loss-of-function mutations are homozygous-lethal (Twell et al., 2002), making it challenging to find an appropriate genetic background in which to express and test the functionality of the transgenic reporter constructs. We produced stable transformants of our two reporter constructs in three different genetic backgrounds, including *Arabidopsis thaliana* Col-0 ecotype (wild type), and two previously uncharacterized *mor1* mutant alleles: *mor1-23* and *mor1-6*. The *mor1-23* allele, which can segregate viable homozygotes, has a T-DNA insertion (SALK_032056) in the final exon 53 (Figure 1A, B) that is retained in the mRNA (Figure 1C). The *mor1-6* allele, generated by EMS mutagenesis in a Col-0 *erecta* background (Till et al., 2003; Torii et al., 1996) (Supplementary Figure 1A), and confirmed to be homozygous-lethal, harbours a single nucleotide mutation at a splice site (G7127A, Figure 1A) that causes retention of intron 25 and a predicted premature stop codon (Figure 1C).

### Expression of MOR1 reporters in the wild-type background is detrimental

When *MOR1-3xYPet* was expressed in the wild-type Col-0 background (Figure 1D) *MOR1* transcript levels were greatly elevated. Quantitative real-time PCR (qPCR) and digital PCR (dPCR) indicated an 8- and 6-fold increase in expression over the wild-type levels, respectively. When the *MOR1-3xYPet* reporter gene was expressed in the *mor1- 23* and *mor1-6* mutant backgrounds, *MOR1* transcript levels were reduced towards wild-type levels (Figure 1D).

We used immunoblotting with a MOR1-specific antibody raised against an N-terminal peptide sequence (Kawamura et al., 2006) (Figure 1B, Supplementary Figure 1B) to analyze protein size and relative abundance in our mutant and transgenic lines. Endogenous wild-type MOR1 produced a 220 kDa band (Figure 1E, lane 1). *mor1-23* homozygotes yielded a weak ∼200 kDa band (Figure 1E, lane 2), consistent with the T-DNA insertion in exon 53 (Figure 1B, 1C) resulting in truncation of MOR1’s C-terminus. Protein extracts from *mor1-6* heterozygotes generated band patterns similar to that of the wild-type extracts (Figure 1E, lane 3), indicating that the *mor1-6* allele (Figure 1B, C) did not yield any visible protein product, which, if present would have an expected molecular weight around 100 kDa.

When the *MOR1-3xYPet* transgene was expressed in the Col-0 wild-type background, endogenous MOR1 (220 kDa) and MOR1-3xYPet (300 kDa) bands were detected with high intensity (Figure 1E, lane 4). Protein extracts from *MOR1-3xYPet* in the *mor1-23* and *mor1-6* backgrounds resulted in immunoblots with the same band patterns but with weaker intensities (Figure 1E, lanes 5 and 6). Intensities of the immunoblotted MOR1 proteins agreed with *MOR1* dPCR transcript analysis such that *MOR1-3xYPet* expressed in the wild-type background had the highest mRNA and protein levels, followed by the *mor1-23* and the *mor1-6* mutant backgrounds (Figure 1D, 1E).

Despite the use of protease inhibitors, there were apparent degradation products. A weak 125 kDa band and a very faint 160 kDa band were detected in the Col-0, *mor1-23* and *mor1-6* extracts. In addition, the MOR1-3xYPet/*mor1-23* and MOR1-3YPet/*mor1-6* extracts resulted in a 220 kDa band that was not encoded by *mor1-23* and *mor1-6* alleles, consistent with separation of the 3xYPet reporter. Based on the relative intensities of the four bands, it is apparent that degradation of the MOR1-3xYPet or the MOR1 endogenous protein in the MOR1-3xYPet/Col-0 line accounts for the greater relative abundance of the 125 kDa degradation product.

Expressing MOR1-3xYPet in the wild-type Col-0 background resulted in reduced root length (Figure 1F, G), indicating that more than one functional copy of the *MOR1* gene is detrimental. Importantly, the MOR1-3xYPet protein rescued the shorter root lengths (Figure 1F, G) of heterozygous *mor1-6* and homozygous *mor1-23* plants, indicating that these lines approximate the wild type most closely. The ability to maintain a *mor1-6* homozygous lethal mutant background when *MOR1-3xYPet* is expressed (Supplementary Figure 1A) further demonstrated that MOR1-3xYPet is fully functional.

### MOR1-3xYPet redistributes to the full length of microtubules in the presence of endogenous untagged MOR1

Live-cell imaging revealed that the distribution of the MOR1-3xYPet reporter along microtubules differed according to the genetic background. When expressed in the wild-type background, MOR1-3xYPet was distributed along the full length of microtubules, whereas in the *mor1-23* and *mor1-6* mutant backgrounds, MOR1-3xYPet was confined to single puncta (Figure 1H). Microtubule growth rates were slightly but significantly reduced when MOR1-3xYPet was expressed in the wild-type background compared to the *mor1-23* background (Supplementary Figure 1C), but were not significantly different between the *mor1-23* and *mor1-6* backgrounds (Supplementary Figure 1D). These findings suggest that the redistribution of MOR1-3xYPet to the full length of the microtubule is associated with higher overall gene expression (Figure 1D), protein levels (Figure 1E), and impaired seedling development (Figure 1G). Since MOR1-3xYPet in the *mor1-23* background yields brighter punctate signals and is not different from the *mor1-6* background in terms of microtubule dynamics (Supplementary Figure 1D), we used the *mor1-23* background but not *mor1-6* for all quantitative imaging experiments reported.

### MOR1 associates with the growing and shrinking microtubule plus end

Analysis of interphase microtubule arrays of epidermal cells in etiolated hypocotyls confirmed that MOR1 is a plus-end tracking protein. MOR1-3xYPet was consistently located at the growing plus ends of mRFP-TUB6-labelled microtubules (Figure 2A-F; Supplementary Movie S1, S2), concentrated within 1 μm from the plus end (Figure 2B). In contrast, we never detected MOR1-3xYPet at the minus ends, which can be identified by their slow depolymerization rates (Figure 2G; Supplementary Movie S3). MOR1- 3xYPet retained its plus-end association during microtubule shrinkage events (Figure 2E; Supplementary Movie S2). We detected MOR1-3xYPet in 93% of 60 depolymerization events (from 7 cells) and found that it persisted on plus ends during catastrophe and subsequent rescue events (Figure 2E; Supplementary Movie S2). MOR1-3xYPet was also rapidly recruited to shrinking microtubule plus ends that form as a result of microtubule severing (Figure 2H, Supplementary Movie S4), which is commonly observed when microtubules encounter and cross over other microtubules at steep angles (> 45°) (Lindeboom et al., 2013; Wasteneys & Ambrose, 2009; Wightman & Turner, 2007; Zhang et al., 2013). After such severing events, MOR1-3xYPet consistently accumulated on the depolymerizing plus ends but was absent from the newly generated minus ends, which also underwent shrinkage (Figure 2H, Supplementary Movie S4).

**Figure 2.**
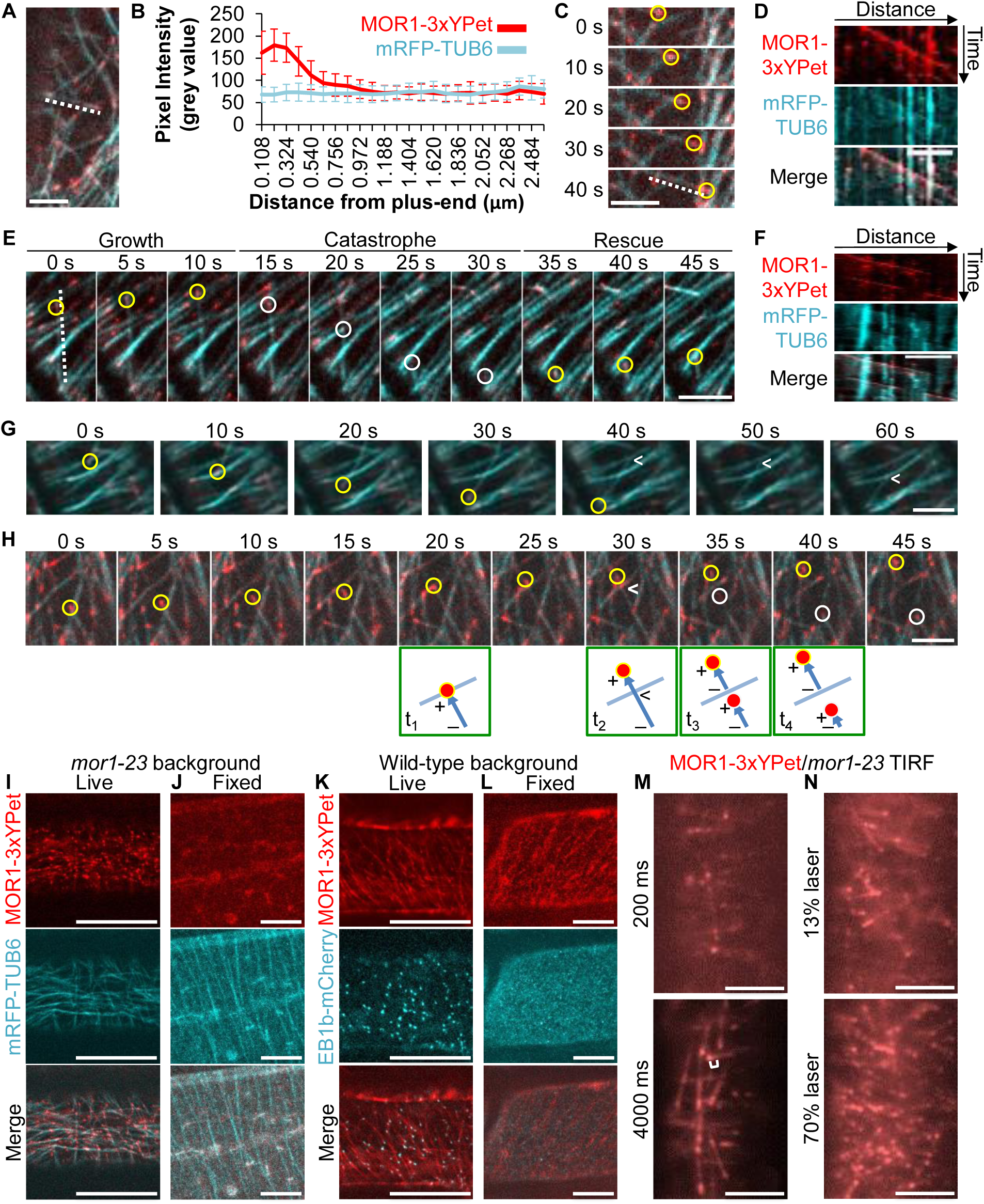
MOR1 binds strongly to microtubule plus ends during microtubule growth, catastrophe and rescue and weakly to the microtubule lattice. **(A)** A spinning-disc confocal image of MOR1-3xYPet (red) on the plus ends of microtubules (cyan). See Supplementary Movie S1. The dotted line marks one microtubule used for fluorescence intensity measurements in **(B)**. Scale bar represents 5 μm. **(B)** Fluorescence intensity plots of MOR1-3xYPet and mRFP-TUB6 line scans of microtubule plus ends. Bars and error bars represent average intensities from 12 line scans ± SD. **(C)** Time-lapse montage of spinning-disc confocal images showing MOR1-3xYPet’s (red) ability to track the plus ends of growing microtubules (cyan). The yellow circles indicate a microtubule plus end. The white dotted line indicates the selection used to generate the kymograph in **(D)**. Scale bar represents 10 μm. See Supplementary Movie S1. **(D)** Kymograph of line selection in **(C)**, collected over 105 s. Scale bar represents 3 μm. **(E)** Time-lapse montage of spinning-disc confocal images of MOR1-3xYPet (red) and mRFP- TUB6 (cyan). MOR1-3xYPet is indicated on microtubule plus end during growth (t = 0-10 s, yellow circle), catastrophe (t = 15-30 s, white circle), and rescue (t = 35-45 s, yellow circle). The dotted line indicates the selection used to create the kymograph in **(F)**. Scale bar represents 5 μm. See Supplementary Movie S2. **(F)** Kymographs of line selection seen in **(E)**, collected over 1 0 s. Scale bar represents 5 μm. **(G)** Time-lapse montage of spinning-disc confocal images of MOR1-3xYPet (red) and mRFP- TUB6 (cyan) showing the polymerizing plus end (yellow circle) and depolymerizing minus end (arrowhead) of a treadmilling microtubule. Scale bar represents 5 μm. See Supplementary Movie S3. **(H)** Time-lapse montage of spinning-disc confocal images of MOR1-3xYPet (red) during microtubule (cyan) crossover and severing. Crossover (t = 25-30 s) is followed by severing (t = 30 s, arrowhead). The newly generated plus end undergoes rapid shrinkage. MOR1-3xYPet is recruited within 5 s (t = 35s, white circle) and tracks the shrinking plus end (t = 35-45 s). MOR1- 3xYPet remains at the growing microtubule’s plus end (yellow circle) throughout. Scale bar represents 5 μm. See Supplementary Movie S4. In the schematic diagram, a growing microtubule (blue) with MOR1 (red circle) crosses over an obstacle microtubule (blue) (t_1_-t_2_). Following crossover (arrowhead), MOR1 tracks the existing, growing plus end as well as the newly generated, shrinking plus end (t_3_-t_4_). + and – indicate the plus and minus ends, respectively. **(I, J)** Confocal imaging of MOR1-3xYPet (red) and mRFP-TUB6 (cyan) in **(I)** live and **(J)** fixed etiolated hypocotyl cells (*mor1-23* background). In live cells, MOR1-3xYPet is plus-end localized. In chemically fixed cells, MOR1-3xYPet is no longer plus-end localized and instead becomes cytoplasmic. Scale bars represent 20 μm. **(K, L)** Confocal imaging of MOR1-3xYPet (red) and EB1b-mCherry (cyan) in **(K)** live and **(L)** fixed cells (wild-type background). In live cells, MOR1-3xYPet is localized along the microtubule lattice, and EB1b-mCherry is plus-end localized; in chemically fixed cells, MOR1-3xYPet no longer appears on microtubule plus ends but still labelled lattice, and EB1b-mCherry is not localized to microtubules and instead becomes cytoplasmic. Scale bars represent 20 μm. **(M, N)** MOR1 is detected in low abundance along the microtubule lattice by altering imaging settings on the TIRF microscope. Live-cell images are taken from the etiolated hypocotyls of *MOR1-3xYPet/mor1-23* seedlings. **(M)** Comparison of images obtained with a 200 ms and 4000 ms exposure time. White brackets indicate the average distance that a single MOR1-3xYPet puncta will travel over a 4000 ms time period. Scale bars represent 5 μm. **(N)** Comparison of images obtained by increasing the laser intensity from 13% to 70% while maintaining the same exposure time. Scale bars represent 5 μm.

### Chemical fixation causes the loss of the microtubule plus end

The plus-end distribution of the MOR1-3xYPet reporter is at odds with previous immunofluorescence results that reported even distribution of MOR1 epitopes along the entire microtubule length (Kawamura et al., 2006). This discrepancy prompted us to determine how chemical fixation, a key step in immunofluorescence experiments, would affect MOR1-3xYPet distribution. After aldehyde fixation, although the general appearance of RFP-TUB6-labelled microtubules remained unchanged, MOR1-3xYPet could no longer be detected on the microtubules (Figure 2I, 2J). This suggested that during chemical fixation, either MOR1-3xYPet’s interaction with microtubules was lost altogether or the plus end disassembled. To distinguish between these two possibilities, we generated a line co-expressing *MOR1-3xYPet* and the plus-end tracking *EB1b- mCherry* (Chan et al., 2003) in the wild-type background, such that MOR1-3xYPet was distributed along the full length of microtubules, and EB1b-mCherry was localized to the plus ends (Figure 2K). Upon fixation, MOR1-3xYPet remained along the full length of microtubules, indicating that the fixation method does not cause MOR1 to dissociate from the microtubule lattice (Figure 2L). In contrast, the EB1b-mCherry signal was lost from the microtubule plus ends but still produced a diffuse cytoplasmic signal (Figure 2K, 2L). These findings indicate that during chemical fixation, the microtubule plus end undergoes partial disassembly, explaining the failure of immunofluorescence to detect MOR1’s preferential association with the microtubule plus end. Nevertheless, a low abundance of MOR1 protein along the microtubule lattice should still be labelled with antibodies resulting in the apparent full-length association. To confirm the presence of MOR1 along the microtubule lattice, we oversaturated the MOR1-3xYPet signal by adjusting exposure time or laser intensity using a Total Internal Reflection Fluorescence (TIRF) Microscope. Despite the strong preference for the plus end, we were able to detect MOR1-3xYPet in low abundance along the lattice (Figure 2M, 2N). We conclude that despite its preferential binding to microtubule plus ends, MOR1 is also capable of microtubule lattice binding, which is enhanced when MOR1 protein level increases in the wild-type background.

### MOR1 associates with all microtubule arrays throughout development and the cell cycle

*MOR1-3xYPet* was expressed in all cell types of root and aerial tissues of seedlings examined (Figure 3A-H), consistent with previous *MOR1* expression analysis (Whittington et al., 2001). In regions of actively dividing cells, MOR1-3xYPet was associated with preprophase bands (PPBs) with strong perinuclear localization (Figure 3I), and with mitotic spindles (Figure 3J, Supplementary Movie S5). During cytokinesis, MOR1-3xYPet was strongly associated with phragmoplast microtubules and enriched at the mid-plane, consistent with microtubule plus-end association (Figure 3K, 3L).

**Figure 3.**
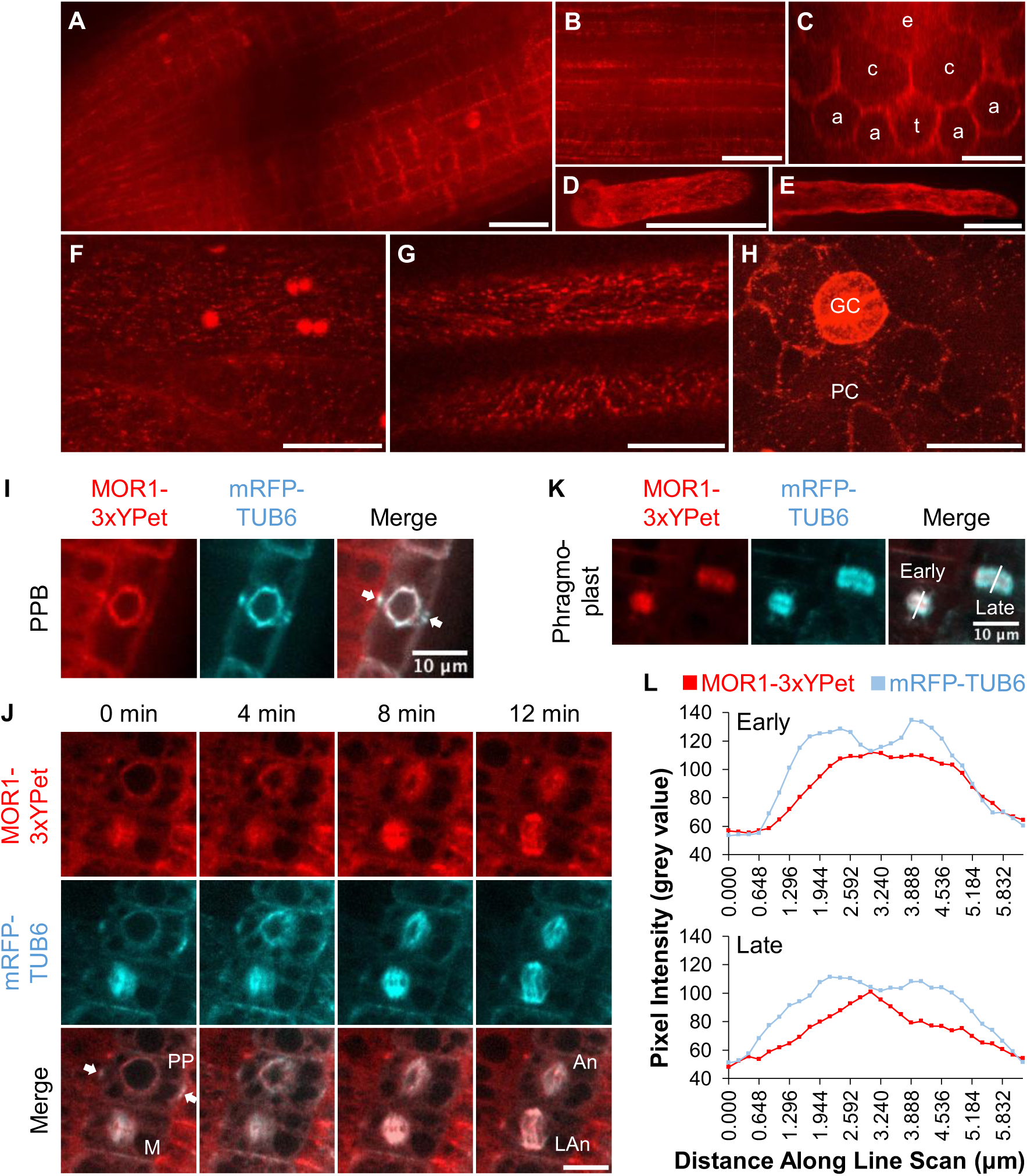
MOR1 localizes to microtubules in various cell types and throughout the cell cycle in *Arabidopsis* seedlings. (A-H) MOR1-3xYPet is expressed throughout *Arabidopsis* seedlings. Scale bars represent 20 μm. **(A)** Root tip cells. The image consists of two stitched images and a merged stack of z-slices taken every 0.5 μm. **(B)** Root elongation zone. Image is a merged stack of z-slices taken every 0.5 μm. **(C)** A cross-section of the root elongation zone showing trichoblasts (t), atrichoblasts (a), cortex (c), and endodermal (e) cells. **(D)** Elongating root hairs. The image is a cortical slice of the root hair. The image is a merged stack of z-slices taken every 0.2 μm. **(E)** Mature root hairs. The image is a cortical slice of the root hair. The image is a merged stack of z-slices taken every 0.2 μm. **(F)** Petiole epidermal cells. The image is a merged stack of z-slices taken every 0.5 μm. **(G)** Etiolated epidermal hypocotyl cells. The image is a merged stack of z-slices taken every 0.5 μm. **(H)** Pavement cells (PC) and guard cells (GC) of cotyledons. The image is a merged stack of z-slices taken every 0.5 μm. **(I-L)** MOR1 is associated with microtubules throughout the cell cycle. **(I)** During pre-prophase, MOR1-3xYPet localizes to microtubules at the preprophase band (PPB, arrows) and around the nuclear envelope as seen in medial sections of a lateral root cap cell. Scale bar represents 10 μm. **(J)** MOR1-3xYPet localizes to microtubules from pre-prophase to late anaphase in dividing root tip cells. Each image shows two cells at different stages of mitosis: the top cell progressed from pre-prophase (PP) to anaphase (An), while the bottom cell progressed from metaphase (M) to late anaphase (LAn). MOR1-3xYPet localizes to microtubules at the PPB (arrows) and around the nuclear envelope during pre-prophase, and re-localizes to spindle microtubules. Scale bar represents 10 μm. See Supplementary Movie S5. **(K)** During telophase, MOR1-3xYPet localizes to the phragmoplast at microtubule plus ends and in microtubule-free domains as seen in root tip cells. Images show two cells at early and late stages of telophase. The middle of the phragmoplast is the newly formed cell plate that excludes microtubules, indicated by a decrease in mRFP-TUB6 signal. The increase in MOR1-3xYPet fluorescence intensity at the middle of the phragmoplast suggests an accumulation of MOR1-3xYPet at microtubule plus ends. **(L)** Fluorescence intensity of line scans of phragmoplasts at early and late telophase shown in **(K)**.

### MOR1’s greater affinity at the plus end is correlated with longer residency

MOR1’s strong association with the microtubule plus end during growth and shrinkage prompted us to develop a method to quantify its relative affinity in living cells. Using a TIRF microscope on variable angle settings equipped with a Fluorescence Recovery After Photobleaching (FRAP) module, we photobleached 10 μm diameter circular regions of interest (ROI) containing several growing microtubule plus ends, then measured the degree of MOR1 recovery by averaging fluorescence within the ROI, while accounting for photobleaching over the course of image acquisition in a similar unbleached ROI, with background fluorescence measured from a third ROI outside of the cell (Supplementary Figure S2). From this, we calculated the mobile fraction (F_m_), the off-rate (koff), and the half-time of recovery (t_1/2_) (Table 1).

**Table 1.** Summary of MOR1 velocities (v) and parameter values (F_m_, k_off_, t_1/2_) of the FRAP model. Mean values ± SE are reported. n_v_ = number of MOR1 particles sampled for velocity calculations, and n_cell_ = number of cells sampled for FRAP experiments.

| Genotype | Treatment | $n_v$ | $v$ ($\mu\text{m min}^{-1}$) | $n_{cell}$ | $F_m$ | $k_{off}$ ($\text{sec}^{-1}$) | $t_{1/2}$ (sec) |
| --- | --- | --- | --- | --- | --- | --- | --- |
| *MOR1-3xYPet/mor1-23* | 21°C | 85 | $7.68 \pm 0.03$ | 31 | $0.75 \pm 0.02$ | $0.28 \pm 0.02$ | $2.78 \pm 0.17$ |
| | 30°C | 113 | $10.49 \pm 0.02$ | 37 | $0.78 \pm 0.02$ | $0.36 \pm 0.02$ | $2.13 \pm 0.11$ |
| *MOR1-3xYPet/mor1-23* | 21°C taxol | 96 | $4.85 \pm 0.15$ | 41 | $0.70 \pm 0.02$ | $0.24 \pm 0.02$ | $3.40 \pm 0.18$ |
| *MOR1-3xYPet/mor1-23/clasp* | 21°C | 87 | $8.70 \pm 0.01$ | 34 | $0.78 \pm 0.02$ | $0.28 \pm 0.02$ | $3.06 \pm 0.23$ |
| *mor1-1-YPet/mor1-23* | 21°C | 60 | $5.03 \pm 0.02$ | 24 | $0.66 \pm 0.03$ | $0.27 \pm 0.02$ | $2.89 \pm 0.21$ |
| | 30°C | 79 | $2.81 \pm 0.02$ | 31 | $0.48 \pm 0.02$ | $0.31 \pm 0.02$ | $2.52 \pm 0.18$ |
| *MOR1-3xYPet/Col-0* | 21°C | 80 | $6.44 \pm 0.03$ | 34 | $0.83 \pm 0.01$ | $0.35 \pm 0.02$ | $2.20 \pm 0.14$ |
| *GFP-CLASP/clasp-1* | 21°C | N/A | N/A | 14 | $0.68 \pm 0.03$ | $0.29 \pm 0.02$ | $2.54 \pm 0.16$ |

We compared MOR1’s affinity for the microtubule plus ends and lattice by quantifying FRAP for MOR1-3xYPet expressed in the *mor1-23* and wild-type backgrounds respectively (Figure 4A, 4B, 4D; Supplementary Movie S6A, S6B). In the *mor1-23* background, the plus-end associated MOR1-3xYPet signal recovered over several seconds (t_1/2_ = 2.78 ± 0.17 sec) with a F_m_ of 0.75 ± 0.02 (Figure 4D; Table 1). In the wild-type background, with MOR1-3xYPet distributed along the full length of microtubules, fluorescence recovered more quickly (t_1/2_ = 2.20 ± 0.14 sec) and the mobile fraction increased (F_m_ = 0.83 ± 0.01) (Figure 4D; Table 1). It is important to note here that the actual turnover of MOR1 on the lattice is probably greater than measured because MOR1’s full-length distribution likely included plus end- as well as lattice-association.

**Figure 4.**
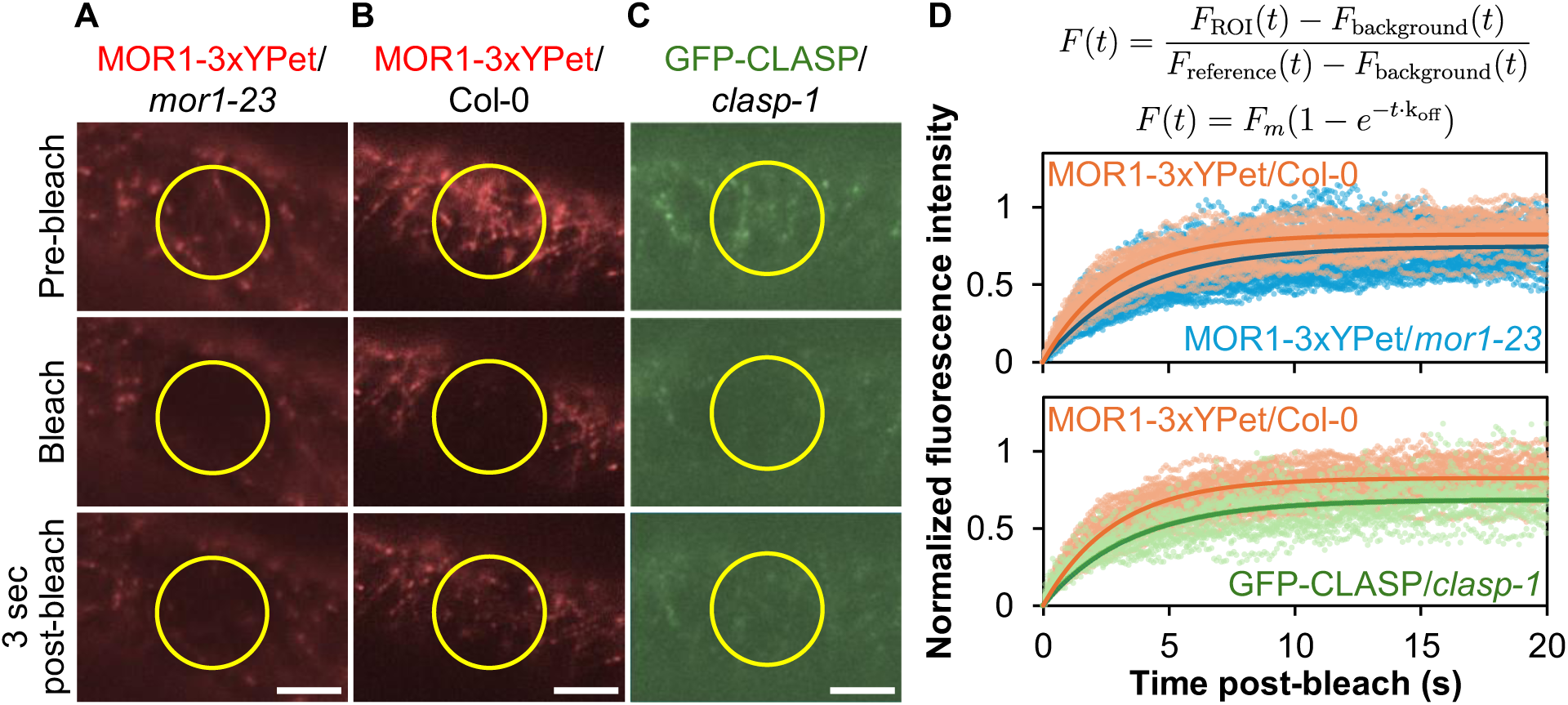
MOR1 displays a stronger association to the microtubule plus end than the lattice. Seedlings Fluorescence recovery after photobleaching (FRAP) experiments were used to measure MOR1 dynamics within a circular region of interest (ROI, yellow circle) in etiolated hypocotyl cells of the following seedlings: **(A)** *MOR1-3xYPet* expressed in the *mor1-23* background, **(B)** *MOR1-3xYPet* expressed in the wild-type Col-0 background, and **(C)** *GFP- CLASP* expressed in the *clasp-1* background as a control for lattice-bound proteins. Scale bars represent 5 µm. **(D)** After subtracting the background signal (F_background_), the ratio of fluorescence intensity within the ROI (F_ROI_) to another unbleached region in the same cell (F_reference_) is used to calculate the mobile fraction (F_m_) and the off-rate (k_off_) of fluorescent-tagged proteins from microtubules.

We were curious to know how MOR1’s lattice-binding coefficients compared to the microtubule-associated protein CLASP, which also has TOG domains but is normally distributed along the length of microtubules (Ambrose et al., 2013). Compared to MOR1- 3xYPet on the lattice, fluorescence recovery was slower for GFP-tagged CLASP (t_1/2_ = 2.54 ± 0.16 sec; Figure 4C, 4D; Table 1) and the mobile fraction was lower (F_m_ = 0.68 ± 0.03), and in fact lower than that measured for plus end-associated MOR1-3xYPet. Together, these results confirm that MOR1’s affinity for the lattice is relatively weak, consistent with its preferred distribution. Despite MOR1’s higher affinity for the microtubule plus end, FRAP analysis indicated that it still undergoes rapid turnover, suggesting that it has limited processivity.

### The mor1-1 L174F mutation in TOG1 reduces polymerization and turnover of MOR1 protein on the microtubule plus end

We next explored the relationship between MOR1’s polymerase activity and its turnover at the microtubule plus end. The *mor1-1* L174F substitution in the N-terminal TOG1 domain (Whittington et al., 2001) greatly reduces microtubule growth and shrinkage rates in a temperature-dependent manner (Kawamura & Wasteneys, 2008). Expressing the *MOR1_pro_:mor1-1-YPet* construct in the *mor1-23* (and *mor1-6*) mutant backgrounds, we were able to recapitulate the *mor1-1* temperature-dependent reduction in microtubule polymerization. For the wild-type *MOR1-3xYPet mor1-23* line, shifting the temperature from 21°C to 30°C increased the microtubule growth rate from 7.68 ± 0.03 μm min^-1^ to 10.49 ± 0.02 μm min^-1^ (Figure 5A, 5B; Table 1; Supplementary Movie S6B, S6C). For the *mor1-1-YPet mor1-23* line, the same temperature shift caused microtubule growth rates to fall from 5.03 ± 0.02 to 2.81 ± 0.02 μm min^-1^ (Figure 5A, 5B; Table 1; Supplementary Movie S6D, S6E).

**Figure 5.**
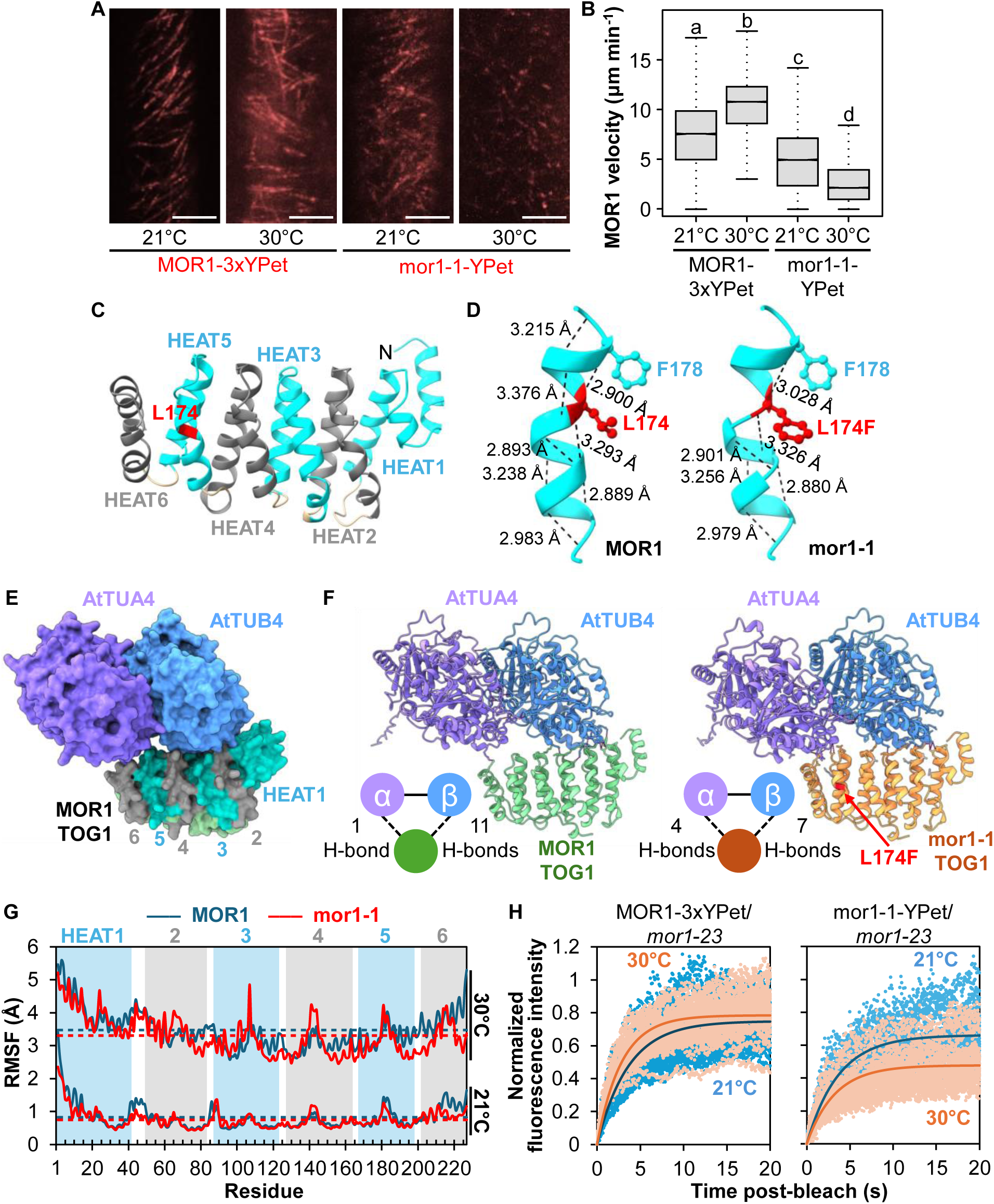
The mor1-1 L174F substitution reduces the rate of microtubule polymerization, alters TOG1 hydrogen bonding and flexibility, and reduces MOR1 turnover on the microtubule plus end. **(A)** Time projections of hypocotyl cells expressing MOR1-3xYPet and mor1-1-YPet at 21°C and 30 C, collected over 44 seconds. Scale bars represent 5 μm. **(B)** Boxplots showing MOR1-3xYPet and mor1-1-YPet velocities at 21°C and 30°C. Horizontal lines mark the median, boxes mark the range from the 25th to 75th percentile, and the whiskers show the highest and lowest data points within 1.5 times the interquartile range. Letters represent statistical differences between genotypes assessed by Kruskal-Wallis and Dunn’s test. n = 85 for MOR1-3xYPet 21°C, n = 113 for MOR1-3xYPet 30°C, n = 60 for mor1-1-YPet 21°C, n = 79 for mor1-1-YPet 30°C. **(C)** Structure of MOR1 TOG1 with six HEAT repeats coloured cyan or grey. The red colour marks the position of the L174 residue. **(D)** The mor1-1 L174F (in red) substitution alters hydrogen bonds within the alpha helix backbone in the fifth HEAT repeat of TOG1. Hydrogen bond lengths (in angstrom Å) are average values from five AlphaFold models. A nearby aromatic residue F178 is also shown. **(E)** An AlphaFold model of MOR1 TOG1 interacting with the *Arabidopsis* α/β-tubulin dimer TUA4 and TUB4. HEAT repeats 1-6 are coloured cyan or grey. **(F)** The mor1-1 L174F substitution (in red) alters the number of hydrogen bonds between MOR1 TOG1 and each tubulin monomer, calculated with a 0.4 Å tolerance. **(G)** Root mean square fluctuation (RMSF, in angstrom Å) of the TOG1 domain of MOR1 and mor1-1, modelled at 21°C and 30°C. Horizontal dashed lines represent mean RMSF values across residues 1-227. **(H)** FRAP data of MOR1-3xYPet and mor1-1-YPet at 21°C and 30°C.

To gain insight into why the mor1-1 L174F substitution reduces microtubule dynamicity, we carried out structural modelling with AlphaFold3 (Abramson et al., 2024). The phenylalanine substitution is predicted to distort one of the antiparallel alpha helices comprising the fifth of six HEAT repeats of the N-terminal TOG domain by altering the number and lengths of hydrogen bonds in the backbone (Figure 5C, 5D). Using Alphafold3 to assess the TOG1 domain’s interaction with tubulin dimers (Figure 5E-G), we found that the mor1-1 L174F mutation caused a more equal dispersion of H-bonds between TOG1 and the two tubulin monomers. The wild-type TOG1 is predicted to have 1 H-bond to ɑ-tubulin and 11 with β-tubulin at HEAT repeats 1, 2 and 5 (Figure 5F). In contrast, the mor1-1 TOG1 domain is predicted to form 4 H-bonds to ɑ-tubulin and 7 H-bonds to β-tubulin (Figure 5F), resulting in a more uniform bond with both tubulins rather than binding predominantly to one. Root mean square fluctuation (RMSF) of the TOG1 domain at 21°C and 30°C indicated that the *mor1-1* mutation decreased the overall flexibility of the entire TOG1 domain, and that this was more pronounced at the restrictive temperature (Figure 5G). In the wild-type TOG1 domain, the minimal binding to α-tubulin could provide the necessary flexibility for efficient dimer detachment, facilitating normal growth rates of the microtubule. This, combined with the overall increase in TOG1 rigidity observed in the RMSF data, could explain the decreased rate of microtubule polymerization in the *mor1-1* mutant at restrictive temperatures, and why there is still a minor but significant reduction in microtubule dynamicity at permissive temperatures.

FRAP results were consistent with mor1-1-YPet having decreased turnover from microtubules at the restrictive temperature (Figure 5H, Table 1). There was a large reduction in mobile fraction for mor1-1-YPet from 0.66 ± 0.03 at 21°C to 0.48 ± 0.02 at 30°C, while there was a slight increase in mobile fraction for the wild type (0.75 ± 0.02 at 21°C to 0.78 ± 0.02 at 30°C). Thus, the *mor1-1* L174F point mutation delays MOR1’s turnover from microtubules, and this is associated with reduced rates of microtubule growth and shrinkage.

### MOR1 turnover from the microtubule plus end is tightly coupled to the microtubule polymerization rate

The correlation between microtubule growth rate and MOR1 turnover suggested that MOR1 residency is strictly coupled with a set number of polymerase reactions. This prompted us to determine if manipulating microtubule dynamics in ways independent of MOR1’s polymerase activity would generate similar effects. We applied 20 μM of the microtubule-stabilizing drug taxol for 4 hours at 21°C. This moderate treatment reduced the F_m_ to 0.70 ± 0.02 and reduced the microtubule growth rate to 4.85 ± 0.15 μm min^-1^ (Figure 6A, 6B, 6E, Table 1; Supplementary Movie S6F). We used CRISPR/Cas9 to knock out the microtubule rescue factor CLASP in the *MOR1-3xYPet*/*mor1-23* line, which resulted in a single nucleotide insertion in exon 6 (Figure 6D), causing a frame- shift, and generated a dwarf phenotype equivalent to the *clasp-1* T-DNA insertion allele (Ambrose et al., 2007). The *clasp* mutation resulted in a higher F_m_ of 0.78 ± 0.02, and faster microtubule growth rate (8.70 ± 0.01 μm min^-1^) (Figure 6C, 6F, Table 1; Supplementary Movie S6G). Similar increases in microtubule growth rates were measured in *clasp-1* mutants (Lindeboom et al., 2019).

**Figure 6.**
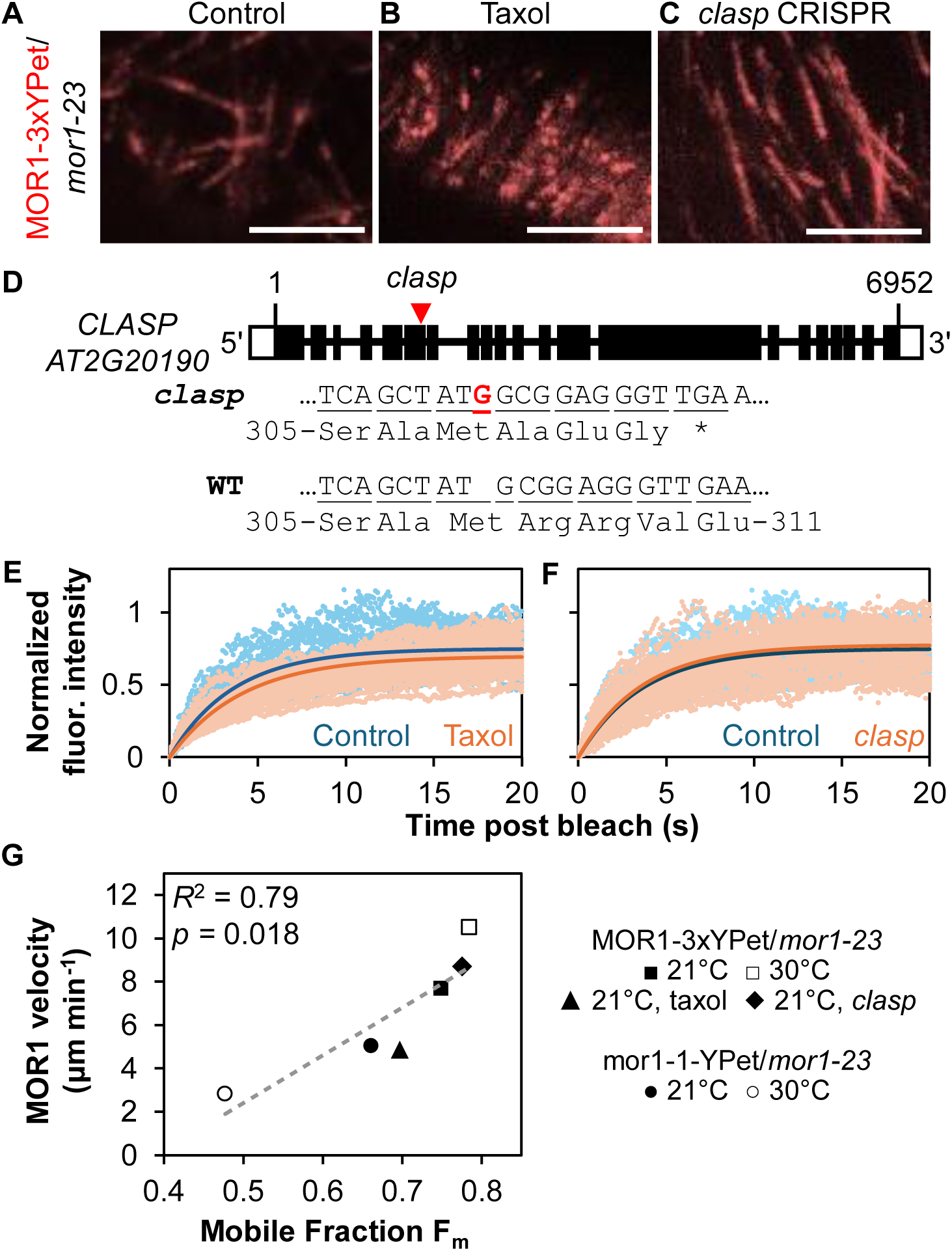
MOR1 turnover at the plus end is tightly coupled to MOR1 velocity and microtubule growth. **(A)** CRISPR/Cas9 generated a single base pair insertion (in red) in exon 6 of the *Arabidopsis CLASP* gene, resulting in a frameshift and a premature stop codon (*). **(B-G)** Microtubule growth is correlated with MOR1 turnover under various experimental conditions. Time projection over 20 seconds taken in etiolated hypocotyl cells expressing: **(B)** MOR1-3xYPet measured at 21°C as the control, **(C)** MOR1-3xYPet under taxol treatment, and **(D)** MOR1-3xYPet in the CRISPR *clasp* background. Scale bars represent 5 µm. All seedlings were also in the *mor1-23* mutant background. **(E)** FRAP data of MOR1-3xYPet from control and taxol-treated cells. **(F)** FRAP data of MOR1-3xYPet from control and CRISPR *clasp* cells. **(G)** MOR1 mobile fraction is positively correlated with MOR1 velocity. Pearson *R*^2^ = 0.79, *p* = 0.018.

Finally, we plotted the mobile fraction against microtubule growth rates for MOR1- 3xYPet and mor1-1-YPet at both 21°C and 30°C, and for MOR1-3xYPet under taxol treatment or in the CRISPR *clasp* mutant background. We found a positive and linear correlation (R^2^ = 0.79) between MOR1 mobile fraction and the MOR1 velocity, a proxy for microtubule polymerization rate (Figure 6G). Under conditions that increase microtubule polymerization, such as higher temperature for wild-type MOR1, the mobile fraction was elevated. Under conditions that reduce microtubule growth rates, such as mor1-1-YPet at 30°C or taxol treatments, the mobile fraction was reduced. Given that our perturbations to microtubule dynamics were both dependent and independent of MOR1’s polymerase activity, the results strongly suggest that MOR1’s turnover is dictated by a finite processivity linked to a fixed number of tubulin subunit additions.

## DISCUSSION

### Microtubule plus end polymerization is coupled to MOR1’s finite processivity

In this study, the application of FRAP to measure the turnover of fluorescently tagged MOR1 on dynamic cortical microtubules has provided new insight into the relationship between MOR1’s polymerase activity and its turnover at the microtubule plus end. Plant cells are ideal for such analysis – their large volume ensures that only a tiny fraction of the fluorescent protein population will be photobleached, and the 2-dimensional nature of the cortical microtubule array enables fluorescence tracking to be recorded within a narrow focal plane close to the plasma membrane. The use of near-TIRF settings further optimized the experimental system by eliminating interference from signals outside the optical section, providing high resolution images for our quantitative analysis.

Our FRAP experiments revealed a strong linear relationship between the rate of MOR1 turnover on the microtubule plus end and the rate of microtubule polymerization. For the wild-type MOR1-3xYPet reporter, increasing microtubule growth rate by shifting the temperature from 21 to 30°C or by knocking out the expression of *CLASP* resulted in more rapid turnover of MOR1 (*i.e.*, reduced residency) whereas taxol treatment, which reduced microtubule growth rate, resulted in slower MOR1 turnover. For the mor1-1- YPet reporter, increasing the temperature from 21 to 30°C greatly reduced both microtubule growth rate and MOR1 turnover. We also found that MOR1 dissociates more quickly when not at the plus end, indicating that its ability to recruit and add tubulin to the microtubule polymer increases its affinity and residency time.

Despite its greater affinity for the plus end, MOR1’s *in vivo* residency is still remarkably brief, given that signal recovery occurs within seconds (t_1/2_ < 3.5 seconds, Table 1). This is remarkably similar to the average XMAP215-GFP *in vitro* residence time of 3.8 seconds (Brouhard et al., 2008). With these findings, we propose that when MOR1 encounters the microtubule plus end, it is drawn into carrying out an obligate but finite series of polymerase reactions.

The strong plus-end preference of microtubule polymerases can be explained by their involvement in the chain reaction of tubulin polymerization. It has been suggested that plus-end tracking proteins recognize specific biochemical or structural characteristics of microtubule plus ends, such as the GTP cap (Zanic et al., 2009), specific tubulin residues, the microtubule seam, or protofilament structures (Akhmanova & Steinmetz, 2008; Carvalho et al., 2003; Sandblad et al., 2006). Given that MOR1-3xYPet also accumulates on the plus ends of depolymerizing microtubules, which comprise GDP-tubulin, the possibility that MOR1 only recognizes the GTP cap can be ruled out. Instead, if encounters with the plus end draw MOR1 proteins into a finite series of polymerase (or depolymerase) reactions, the relative dwell time will be greater than with encounters along the lattice.

An obvious question that arises is how finite processivity can be achieved. It was previously speculated that binding of a tubulin heterodimer to the N-terminal TOG1 domain could initiate a conformational change in the C-terminal domains of MOR1 such that the protein partially dissociates and “inchworms” forward along the same protofilament track upon completion of each polymerization event to enable addition of a tubulin dimer (Kawamura & Wasteneys, 2008). A ratchet-like mechanism for adding subunits was suggested by evidence that the TOG1 domain of yeast Stu2 preferentially binds to curved αβ-tubulin and has less affinity for straight, polymerized tubulin (Ayaz et al., 2012). Both models predict that MOR1 and its homologues work processively to add tubulin subunits with minimal dissociation. Our current data do not support either model; we showed that MOR1 puncta recovered within seconds of photobleaching, suggesting that MOR1 undergoes complete dissociation from the microtubule after incorporating a set number of tubulin subunits. One compelling explanation for finite processivity is a sidewinder model (Figure 7) whereby MOR1 switches tracks to the adjacent protofilament after each polymerization event, but eventually encounters the microtubule seam, a barrier for track switching that leads to MOR1’s dissociation.

**Figure 7.**
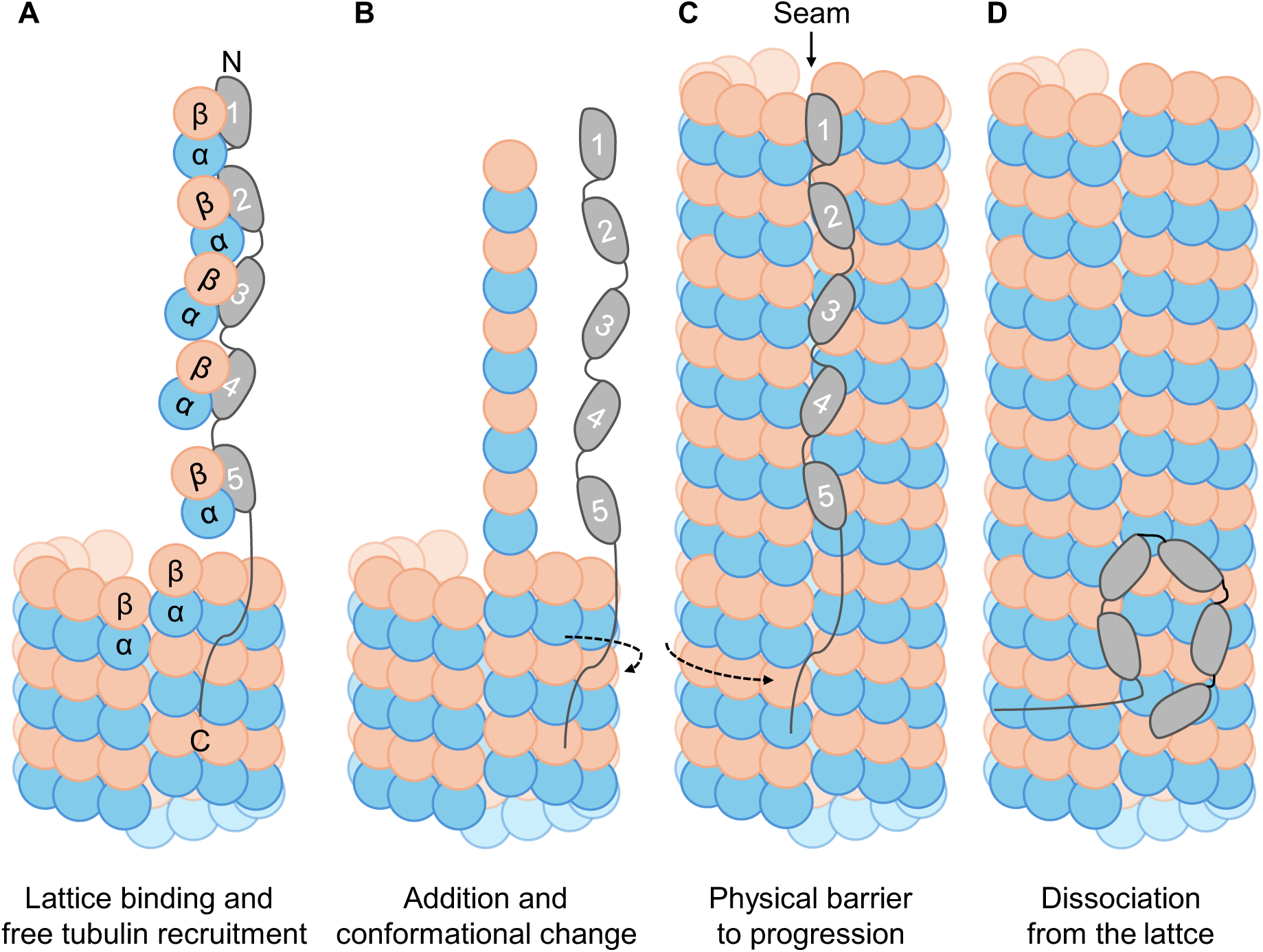
A sidewinder model of MOR1’s polymerase activity and finite processivity at the microtubule plus end. **(A)** MOR1 C-terminus binds the microtubule lattice. TOG domains recruit free tubulin dimers. **(B)** TOG domains add free tubulin dimers to the plus end and detach from the dimers. This process alters MOR1’s conformation such that MOR1 positions itself in the next protofilament. MOR1 continues to incorporate free tubulin into the next protofilament. If MOR1 fails to incorporate tubulin dimers (such as the case for lattice-bound MOR1), MOR1 will dissociate. If MOR1 has impaired polymerase function (such as the case for mor1-1), it will take longer to add dimers and move more slowly from one protofilament to the next. **(C)** MOR1 moves across all 13 protofilaments and encounters a microtubule seam where α/β-tubulin units do not line up perfectly. **(D)** MOR1 may cross the seam to continue polymerization or dissociate from the microtubule.

According to the sidewinder model, each MOR1 could add 13 tubulin dimers during one rotation around the microtubule plus end – assuming only one tubulin dimer is added for each polymerase reaction – or a multiple of 13 if two or more TOG domains act as recruitment templates. Brouhard et al. (2008) estimated under *in vitro* conditions that each XMAP215 protein catalyzes the addition of at least 25 tubulin subunits to growing microtubules (Brouhard et al., 2008), so it seems unlikely that just one tubulin dimer is added. To determine the number of tubulin dimer additions per MOR1 residency at the plus end, we used 1/k_off_ to estimate how long each MOR1 protein stayed on the microtubule, and then determined how many tubulin dimers were added over this time. With the assumptions that 13 MOR1 proteins could occupy the plus end, and that each tubulin dimer is ∼8 nm in length, we estimate that each MOR1-3xYPet adds approximately 60 tubulin dimers per encounter, with 58 tubulins at 21°C and 61 at 30°C (see methods for details). Interestingly, in the absence of CLASP, which might otherwise compete with MOR1 for the microtubule plus end and/or increase the free tubulin concentration, the estimate is 65 tubulins, which would suggest that all five TOG domains could be adding tubulin dimers to each protofilament, as previously suggested for the 5-TOG XMAP215 (Kerssemakers et al., 2006). Similarly, the Polarized Unfurling Model – based on cryo-electron microscopy images of the 2-TOG Alp14 homodimers complexing with free tubulin *in vitro* – proposes that the 4-TOG homodimer Alp14 adds 4 tubulins to each protofilament with each reaction (Nithianantham et al., 2018). Further refinement of our FRAP system, however, will be required to get more accurate estimates. We discuss these limitations in a separate section below.

What is the mechanism that accounts for the increased turnover when MOR1 is lattice-bound? An *in vitro* study by Geyer et al. (2018) showed that TOG-tubulin interactions largely eliminate lattice binding in both Stu2 from budding yeast and Zyg-9 from *Caenorhabditis elegans,* consistent with plus-end preference being linked to polymerase activity. Lattice-bound MOR1 should still be able to associate with free tubulin through TOG1, but considering the near impossibility for tubulin to be incorporated along the lattice, the protein most likely fails to undergo a conformational change, and thus dissociates entirely. By contrast, we found that the turnover of CLASP protein was much slower. CLASP normally binds along the length of cortical microtubules, where, in addition to its role as a rescue factor (Ambrose et al., 2011), it fosters the recycling of the auxin transporter PIN2 and the brassinosteroid receptor BRI1 through a direct interaction with the retromer complex associated protein sorting nexin 1 (Ambrose et al., 2013; Ruan et al., 2018). Its longer residency is consistent with these additional cellular functions.

### Insights into the mor1-1 L174F phenotype

The MOR1 gene was originally identified in a mutant screen that identified the temperature-sensitive *mor1-1* mutant (Kawamura et al., 2006; Kawamura & Wasteneys, 2008; Whittington et al., 2001). Subsequent studies determined that the substitution of leucine 174 in for phenylalanine in the N-terminal TOG domain (TOG1) greatly reduces microtubule growth rates at the restrictive temperature (Kawamura & Wasteneys, 2008). This, in turn, reduces the frequency of microtubule encounters, resulting in loss of cortical microtubule self organization as confirmed by mathematical modelling (Allard et al., 2010), and a profound loss of growth anisotropy through altered cell wall mechano- chemical properties (Fujita et al., 2011). Our current study provides a compelling explanation for the *mor1-1* phenotype. AlphaFold3 structural modelling predicted conformational changes in the fifth of six HEAT repeats of TOG1, and associated changes in TOG1’s interaction with tubulin dimers. Specifically, the mor1-1 TOG1 domain is predicted to have a reduced flexibility, with increased H-bonds with α-tubulin, and fewer with β-tubulin. These predicted structural changes help to explain why the mor1-1 protein works more slowly to catalyze tubulin addition, and why its turnover from the microtubule plus end is delayed. These findings are consistent with a greatly reduced dissociation constant for recombinant MOR1 TOG1-TOG2 with the L174F substitution when co-sedimented with taxol-stabilized microtubules (Lechner et al., 2012). Interestingly, size exclusion chromatography measurements in the same study indicated that the L174F substitution did not affect binding to free tubulin (Lechner et al., 2012). Thus, the mor1-1 protein is more likely to be delayed in dissociation after tubulin polymerization is complete.

### Limitations of using FRAP to infer MOR1-microtubule interactions

Our FRAP technique is limited in that k_off_ does not perfectly reflect MOR1 turnover on the microtubules immediately after photobleaching. This is because a stream of unbleached cytoplasmic MOR1 quickly moves into the ROI, potentially obscuring MOR1 turnover on the plus end. Since k_off_ is likely overestimated due to the rapid recovery of cytoplasmic signal after photobleaching, MOR1 residency time and the amount of tubulin addition are likely underestimated. This effect will be more pronounced if the plus end signal is dim or if turnover on the plus end is very slow (such as mor1-1-YPet at 30°C) such that cytoplasmic signal recovery accounts for a high proportion of recovery immediately after photobleaching. While it is possible to measure only signal recovery on the plus end, data processing is more challenging due to MOR1 particles moving in and out of focus during the recovery period, so we opted to capture average signals over a larger area and depth. Under this constraint, the mobile fraction was found to be a better predictor of MOR1 turnover because after 20 seconds of recovery, cytoplasmic streaming has likely redistributed unbleached cytoplasmic MOR1 to the pre-bleached state. While the mobile fraction represents the re-association of unbleached MOR1 to the plus end, the immobile fraction could either represent a population of MOR1 with higher residency time or the re-association of bleached MOR1 to the plus end.

### MOR1 is a plus-end tracking microtubule polymerase and depolymerase

By successfully generating translational MOR1 fluorescent reporters for live-cell imaging, we determined that MOR1 has a strong association with both polymerizing and depolymerizing microtubule plus ends. This supports MOR1’s dual function as a microtubule polymerase and depolymerase, as earlier suggested by the i*n vivo* reduction in microtubule growth and shrinkage rates in the *mor1-1* mutant at restrictive temperature (Kawamura & Wasteneys, 2008). This finding is consistent with *in vitro* evidence that XMAP215-GFP tracks both the growing and shrinking plus ends of microtubules (Brouhard et al., 2008), and also with earlier in vitro studies that XMAP215 could increase the rate of microtubule shortening velocity (Vasquez et al., 1994), or to act as a microtubule-destabilizing factor (Shirasu-Hiza et al., 2003). Bridging *in vitro* and *in vivo* data is important for elucidating MOR1’s function due to differences in microtubule dynamics in the two systems. For instance, microtubule shrinkage *in vitro* is achieved by removing tubulin from the reaction solution (Brouhard et al., 2008), whereas *in vivo*, tubulin levels are relatively constant, and only some microtubules within a population undergo shrinkage at one instant under steady state conditions. By working in reverse, microtubule polymerases might coordinate a form of controlled depolymerization, thereby fostering rescue instead of complete catastrophe.

Although MOR1’s strong preference for the microtubule plus end is expected, previous immunofluorescence analysis (Kawamura et al., 2006) indicated that MOR1 was evenly distributed along the full length of microtubules, in contrast to its homologues XMAP215 and Stu2 (Brouhard et al., 2008; Podolski et al., 2014). We have now resolved this discrepancy by showing that aldehyde fixation, through failure to preserve the dynamic microtubule plus end, eradicates plus-end-specific MOR1-3xYPet and EB1b-mCherry signals, but does not cause MOR1-3xYPet to dissociate from the lattice. This finding brings into question many previous immunofluorescence studies on plus-end tracking proteins that will require further validation from live-cell imaging.

### MOR1 copy number and expression is tightly regulated for proper growth and development

When the MOR1-3xYPet construct was expressed with the native MOR1 promoter in the wild-type background, overall MOR1 expression more than doubled, suggesting a positive feedback in MOR1 transcriptional regulation when two copies of wild-type *MOR1* gene are present. This results in high MOR1 protein level and the distribution of MOR1-3xYPet along the full length of microtubules. Since increased microtubule lattice labelling has been demonstrated when plus-end tracking proteins were overexpressed (Carvalho et al., 2003), our previous finding that MOR1 TOG1-TOG2-GFP had a full-length microtubule distribution pattern (Lechner et al., 2012) can be explained by its co-expression with endogenous, full-length MOR1.

As a result of expressing two copies of wild-type *MOR1* and increased MOR1 lattice association, plants showed slower microtubule growth and shorter roots. This suggests that *MOR1* expression needs to be precisely controlled to maintain proper growth and development. Too much MOR1 might have blocked other MAPs, such as CLASP, from interacting with the microtubule lattice, resulting in altered microtubule dynamics. Overall, our findings provide a compelling explanation for why MOR1 in plants, and its homologues in other eukaryotic systems, retain single copy gene status; the detrimental phenotypic effects of having multiple copies expressed could provide strong selective pressure for the attrition of extra copies.

### MOR1 may be functionally distinct from its homologues

Our findings establish MOR1’s role as a microtubule polymerase and depolymerase with a finite processivity. MOR1’s involvement in both microtubule growth and shrinkage raises the possibility that MOR1 turnover is regulated by catastrophe factors such as the ARMADILLO-REPEAT KINESIN 1 (Eng & Wasteneys, 2014) at the plus end. In addition to its polymerase activity, MOR1’s rapid recruitment to the depolymerizing plus ends following microtubule crossover suggests that MOR1 also participates in microtubule depolymerization following KATANIN-mediated severing (Lindeboom et al., 2013; Wightman & Turner, 2007; Zhang et al., 2013). It remains unclear if MOR1 interacts with other MAPs to control microtubule dynamics.

A better understanding of MOR1 can inform functional diversity of proteins in the XMAP215/Dis1 family. Although we can draw functional comparisons to other systems, it is important to point out that most studies on MOR1 homologues were conducted *in vitro,* and that fungal homologues Stu2 and Alp14 have two TOG domains and a distinct C-terminal structure compared to MOR1 (Al-Bassam & Chang, 2011). While fission yeast Alp14 was noted to promote both microtubule plus- and minus-end polymerization (Al-Bassam et al., 2012; Podolski et al., 2014), MOR1 is not found at the minus end. Recently, XMAP215 (Thawani et al., 2018), Stu2 (Gunzelmann et al., 2018), and Alp14 (Flor-Parra et al., 2018) were noted to play a role in microtubule nucleation in conjunction with the gamma-tubulin ring complexes. Further studies are required to determine if MOR1 plays similar roles in controlling minus end dynamics or nucleating microtubules.

### Concluding remarks

Our study provides insight into the properties of MOR1 as a microtubule polymerase and depolymerase, and offers several technical advancements for research on MAPs. With the development of our FRAP method, we can begin to apply this technique to evaluate the role of other MAPs in the regulation of cytoskeletal dynamics.

## MATERIALS AND METHODS

### Generation of transgenic plant materials

The *MOR1_pro_:MOR1-3xYPet* and *MOR1_pro_:mor1-1-YPet* constructs were generated through recombineering. The *MOR1* gene (*AT2G35630*) in the TAC clone JAtY62C20 was tagged with either *YPet-6xHis* or *3xYPet-6xHis* using the primers At2g35630F and At2g35630R and the Universal *YPet* or *3xYPet* recombineering cassettes as templates following previously described procedures described (Alonso & Stepanova, 2014; Zhou et al., 2011). The insertion of the tags was confirmed by PCR and sequencing using the primers *At2g35630*TestF and *At2g35630*TestR. Next, the 67 kb insert in the JAtY62C20 clone was trimmed by recombineering using Ampicillin and Tetracycline cassettes and the primer pairs *At2g35630*LBdel + replaRB-amp and *At2g35630*RBDel + replaLB-tet, respectively, leaving the *MOR1* genomic sequence plus 11.5kb upstream of the ATG and 5.3 kb downstream of the stop codon. These recombination events were confirmed using the primer pairs *At2g35630*TestDelLB + RBTest and *At2g35630*TestDelRB + LBTest. To introduce the *mor1-1* mutation in the tagged clones, we first inserted the *GalK* selectable marker gene using the *GalK* recombineering cassette (Zhou et al., 2011) using the primers *mor1-1*GalKF and *mor1-1*GalKR. The *GalK* cassette was then replaced by a DNA fragment containing the *mor1-1* mutation using genomic DNA as template and the primers *mor1-1*repF and *mor1-1*repR. The introduction of the *mor1-1* point mutation was confirmed by sequencing using the primers *mor1-1*testF and *mor1- 1*testR. Primers used for recombineering are listed in Supplementary Table S1.

The recombineered constructs *MOR1_pro_:MOR1-3xYPet* and *MOR1_pro_:mor1-1-YPet* were miniprepped and transferred into *Agrobacterium tumefaciens,* which was used to transfect the *Arabidopsis* heterozygous SALK_032056 T-DNA insertion line (herein referred to as *mor1-23*) using a floral dipping technique for transforming large TAC constructs (Alonso & Stepanova, 2014). After selection of T1 lines by BASTA resistance, T2 seedlings homozygous for each transgene (BASTA resistance) were also screened for either homozygous wild-type *MOR1* or *mor1-23* backgrounds. These seedlings were taken to T3 and subsequent generations to be used for experiments. To obtain transgenic reporters in the *mor1-6* homozygous-lethal background, we crossed *MOR1_pro_:MOR1-3xYPet* and *MOR1_pro_:mor1-1-YPet* plants in the homozygous *mor1-23* background with *mor1-6* heterozygotes (which was also homozygous for *erecta105*), and selected lines homozygous for the transgene reporters by BASTA resistance and the *mor1-6* allele by sequencing in the F3 generation. Primers used to genotype *mor1- 23*, *mor1-6*, and *erecta105* backgrounds are listed in Supplementary Table S1.

To visualize the MOR1 reporters and microtubules simultaneously, homozygous T3 lines of *MOR1_pro_:MOR1-3xYPet* and *MOR1_pro_:mor1-1-YPet* were transfected with a *35S_pro_:mRFP-TUB6* construct (provided by Dr. Richard Cyr, Pennsylvania State University) and homozygous T3 lines were used for further experiments. T3 lines of *MOR1_pro_:MOR1-3xYPet* in the *mor1-23* background were also crossed into a *EB1bpro:EB1b-mCherry* reporter line (provided by Dr. Ram Dixit, Washington University St Louis) and F3 lines (homozygous for the *MOR1_pro_:MOR1-3xYPet* and *EB1b_pro_:EB1b- mCherry* transgenes, and for the *mor1-23* T-DNA insertion) were used for further experiments. Col-0 lines were used for all wild-type plants.

To generate a *clasp* mutant background in the *MOR1_pro_:MOR1-3xYPet* line, an egg cell-specific promoter was used to express guide RNAs (gRNAs) targeting the *CLASP* gene (*AT2G20190*) for CRISPR/Cas9 (Wang et al., 2015). Briefly, two 20 bp gRNA targets were found with ChopChop (https://chopchop.cbu.uib.no) (Labun et al., 2016). The target sites were PCR amplified with primers containing gRNA target sequences using the pCBC-DT1T2 vector as a template (Addgene plasmid # 50590; http://n2t.net/addgene:50590; RRID:Addgene_50590). The resulting 626 bp band was cut from an agarose gel and purified. The fragments were then ligated into the pKEE401 plasmid (Addgene plasmid #91715; http://n2t.net/addgene:91715; RRID:Addgene_91715) using BsaI (NEB) and T4 ligase (NEB). The ligated plasmid was transformed into Subcloning Efficiency DH5α competent cells (Invitrogen), miniprepped, sequenced, transformed into *Agrobacterium* GV3101 (GoldBio), and introduced into *MOR1-3xYPet/mor1-23* lines via floral dip (Clough & Bent, 1998). Transformants were selected with kanamycin and sequenced to confirm the presence of the *CLASP* point mutation. T3 seedlings homozygous for the *clasp* point mutation were used for FRAP experiments. Primers used for CRISPR are listed in Supplementary Table S1.

### Plant growth conditions

Seeds were sterilized with ethanol and hydrogen peroxide, resuspended in water, and plated onto Petri dishes containing half-strength Murashige-Skoog media (Phytotech Labs) and 1.2% Bacto Agar (BD Diagnostics) with no sucrose. The plated seeds were stored in the dark at 4°C for 2-3 days before being transferred to a 21°C growth cabinet with 24 h light. For experiments using dark grown hypocotyls, the plates were wrapped in foil (to stimulate etiolation) and then transferred to a 21°C growth cabinet for 4 days. For experiments conducted at 30°C, the plates were moved to a 30°C growth chamber for 24 h prior to imaging.

### Chemical fixation of seedlings

Six-day-old seedlings grown in the dark were fixed in PME buffer (25 mM PIPES, 0.5 mM MgSO_4_, 2.5 mM EGTA, pH 7.2) with 0.5% glutaraldehyde and 1.5% paraformaldehyde for 30 minutes under vacuum infiltration. Seedlings were then washed three times with PMET buffer (PME buffer with 0.05% Triton-X 100) before being placed under a cover slip for imaging.

### RNA extraction, reverse transcription-PCR (RT-PCR), and quantitative real-time PCR (qRT-PCR)

Six-day-old seedlings were used for gene expression analysis. About 30 seedlings from each genotype were ground in the TRIZOL reagent (Invitrogen, Life Technologies) for RNA extraction. The quality of the RNA was assessed on an agarose gel before proceeding. Samples were treated with DNase I (Amplification Grade, Invitrogen) to remove any residual genomic DNA, and then subjected to reverse transcription with SuperScript III Reverse Transcriptase (Invitrogen, Life Technologies) to obtain complementary DNA (cDNA). For RT-PCR, cDNA templates were used directly in a PCR mix and amplified with primers flanking *mor1-23* and *mor1-6*. For qRT-PCR, cDNA templates were amplified with *MOR1*-specific primers and the PowerUp SYBR Green Master Mix (Thermo Fisher Scientific, USA) on an Applied Biosystems™ QuantStudio™ 5 Real-Time PCR machine. *MUSE3* (*AT5G15400*) was used as an internal control for normalization. Primers used for RT-PCR and qRT-PCR are listed in Supplementary Table S1.

### Digital PCR (dPCR)

Digital PCR Master Mix (4X) was made of 8 μL (25X) SYBR Green I Nucleic Acid Gel Stain (Thermo Fisher Scientific, USA) and 80 μL Absolute Q™ DNA Digital PCR Master Mix (5X) with water added to reach a final volume of 100 μL. Each dPCR assay mix contained 2 μL cDNA, 2.5 μL Digital PCR Master Mix (4X), 0.2 μL *MOR1* forward primer (same as qRT-PCR primer) and 0.2 μL *MOR1* reverse primer (same as qRT-PCR primer) with water added to reach a final reaction volume of 10 μL. 8 μL of each assay mix was loaded into the sample well of a QuantStudio™ Absolute Q™ MAP16 Plate Kit (Thermo Fisher Scientific, USA) and 15 μL of QuantStudio Absolute Q Isolation Buffer (Thermo Fisher Scientific, USA) loaded into each of the wells. The plate was then transferred to the QuantStudio Absolute Q Digital PCR System (Thermo Fisher Scientific, USA). dPCR data were analysed using the Applied Biosystems QuantStudio Absolute Q Digital PCR Software (Version: 6.2.1).

### Affinity purification of MOR1 antibodies using synthetic antigens

MOR1 antibodies used by Kawamura et al. (2006) were purified using procedures described by Liu et al. (2009). 2 mg synthetic MOR1 antigen peptide (Cys-βAla- TRKIRSEQDKEPEAE, Kinexus) was dissolved in 1 mL Coupling Buffer (50 mM Tris-HCl, 5 mM Na_2_EDTA, pH 8.5). 2 mL Sulfolink™ resin slurry (Thermo Scientific) was centrifuged for 3 minutes at 14,000 x g. The supernatant was discarded, and the resin was washed six times with Coupling Buffer. 1 mL each of dissolved peptide and washed resin were combined on a rotator for 15 minutes at room temperature, then left stationary for 30 minutes at room temperature. The resin was resuspended, transferred to an empty SulfoLink™ Resin column (Thermo Scientific), and immediately washed with 2 mL Coupling Buffer. Resin slurry and coupled peptide were blocked by adding 1 mL 50 mM cysteine in Coupling Buffer, incubated on a rotator for 15 minutes at room temperature, then left stationary for 30 minutes at room temperature. The gravity-flow column was drained down to the top of the resin bed (without letting the resin bed dry out), washed with 16 mL 1 M NaCl in Coupling Buffer, then washed with 16 mL 0.05% (w/v) sodium azide in 10 mM Tris-HCl pH 7.8 and stored at 4°C.

3 mL rabbit sera terminal bleed containing MOR1 polyclonal antibodies was diluted with 5 mL PBS (10 mM Na_2_HPO_4_, 1.8 mM KH_2_PO_4_, 137 mM NaCl, 2.7 mM KCl, pH 7.4) with 0.01% (w/v) sodium azide and applied to the column on a rotator overnight at 4°C. The column was drained under gravity and washed with 10 mL RIPA buffer (50 mM Tris pH 7.5, 150 mM NaCl, 1% (v/v) nonyl phenoxylpolyethoxyl ethanol (NP-40), 0.5% (w/v) sodium deoxycholate, 0.1% (w/v) SDS), 10 mL lysis buffer (20 mM Tris pH 8.0, 1 M NaCl, 1 mM Na_2_EDTA, 0.5% (v/v) NP-40), and 10 mL 10 mM Tris-HCl pH 7.8. Purified antibody was eluted with 0.5 mL aliquots of 100 mM glycine pH 2.5, into tubes containing 0.5 mL 1 M Tris-HCl pH 7.8. The Bradford Assay (Bio-Rad) was used to determine protein concentrations with optimal values of 50-100 µg. Finally, the column was neutralized with 10 mL 0.05% (w/v) sodium azide in 10 mM Tris-HCl pH 7.8, and stored at 4°C for future use.

### SDS-PAGE and immunoblotting with MOR1 antibodies

Whole seedlings at 10 days after germination were flash-frozen in liquid nitrogen and ground using a mortar and pestle. Protein was extracted in RIPA buffer with added Halt™ protease and phosphatase inhibitor cocktail (Thermo Scientific), filtered through two layers of Miracloth, and centrifuged at 4500 x g for 10 minutes at 4°C. Supernatant was retained, and protein concentration was determined by Pierce™ BCA Protein Assay (Thermo Scientific). Protein lysate was boiled in 4x Laemmli SDS sample buffer (to a final concentration of 62.5 mM Tris-HCl pH 6.8, 2% (w/v) SDS, 10% (v/v) glycerol, 5% (v/v) β-mercaptoethanol, 0.001% (w/v) bromophenol blue) for 10 minutes at 95°C. 30 µg boiled protein lysate was loaded into each of two 4-12% Bis-Tris polyacrylamide gels (Invitrogen) and ran at 100 V in 1x MOPS SDS running buffer (Invitrogen) for 2.5 hours. The first gel was rinsed and transferred to a 0.2 µm PVDF membrane (Bio-Rad) overnight at 4°C with constant current. The membrane was briefly washed in dH_2_O, stained with Ponceau S (Thermo Scientific) to confirm successful protein transfer prior to immunoblotting, imaged, and washed 3 x 1 minute in dH_2_O or until no stain remained. Membrane was blocked with 5% (w/v) bovine serum albumin in TBST buffer (20 mM Tris-HCl pH 7.5, 150 mM NaCl, 0.1% (v/v) Tween 20) for 1 hour, then probed overnight with primary antibody (Anti-MOR1, terminal bleed fraction 4, diluted 1:100 with the blocking solution). Membrane was washed 3 x 10 minutes in TBST, probed with secondary antibody (Anti-Rabbit HRP 1:10000, Sigma) for 1 hour, washed 3 x 10 minutes in TBST, and developed with Supersignal West Atto Ultimate sensitivity substrate (Thermo Scientific). The second gel was stained with Pierce™ Silver Stain or Coomassie Blue (Thermo Scientific) for protein visualization or as a gel loading control. All gels and blots were imaged and analyzed using Chemidoc with the Image Lab software (Bio-Rad).

### Confocal microscopy

Imaging of the live- and fixed-samples was done primarily on a Perkin Elmer Ultraview VoX Spinning Disc Confocal system (Perkin-Elmer) mounted on a Leica DMI6000 B inverted microscope and equipped with a Hamamatsu 9100-02 electron multiplier CCD camera (Hamamatsu). An argon 516 nm laser line with a complementary YFP (540/30) emission band-pass filter or a 561 nm laser with a complementary RFP (595/50) emission band-pass filter was used. 5 to 7-day old seedlings (either light- or dark-grown) were used for imaging. Time-lapse images were acquired every 5 seconds with a 63x (glycerol) objective lens (NA = 1.3). Optical z-stacks of 0.2 – 1.0 μm slices were acquired.

To obtain deeper optical z-stacks of roots, a Leica SP8-FLIMan confocal microscope with a 63x (water) objective lens (NA = 1.2) was used. Seedlings expressing *MOR1- 3xYPet* were placed on coverslips in water and affixed under a 1% agarose pad. Images were acquired at 0.5 μm increments. To excite YFP, a 514 nm laser was used and fluorescence emission was detected by HyD SMD detectors, set to detect 525-560 nm for YFP. A line average of 4 was used for the experiments.

### Variable-Angle TIRF Microscopy

Whole seedlings were mounted in liquid half-strength Murashige-Skoog media (Phytotech Labs) in round glass bottom microwell imaging dishes with the coverslip at the bottom (MatTek Corporation, Part No. P35G-1.5-20-C). Several blocks of agar growth medium were placed atop the sample to ensure the tissue was held close to the coverslip. For experiments conducted at 30°C, all materials including forceps and mounting media were pre-warmed to 30°C.

The TIRF microscopy system was based on a Zeiss Axiovert Z1 microscope, equipped with a Zeiss TIRF III slider, a diode-pumped solid state laser, a Q imaging Rolera EM-C2 EM-CCD camera, and for later experiments a Hamamatsu Orca Flash sCMOS camera, and the Zeiss DirectFRAP imaging system, all controlled by Zen 2 software (blue edition, Zeiss). A 488 nm laser with a 525/50 nm emission filter and a 63X/1.46 Oil Korr TIRF objective lens were used for imaging YFP. The microscope stage, stage cover, and an objective lens heater were controlled by a temperature module (PeCon GmbH) to maintain a constant temperature of either 21°C or 30°C throughout imaging, which was verified by measuring the temperature of the immersion oil with a probe.

For FRAP experiments, the apical part of the hypocotyl (where cells are rapidly expanding) was used for image acquisition. Three pre-bleach images were acquired prior to photobleaching. A circular region of interest (ROI, area = 80.85 μm^2^) was used for photobleaching. The 488 nm laser was set to 100% laser power for 800 ms and the beamsplitter at 50% TIRF/50% Manipulation to bleach the fluorescent proteins within the ROI. Post-bleach images were collected every 100 ms for up to 40 seconds. Five to ten movies were collected per plant and five to ten individual plants were used for each genotype.

### Drug treatment

Paclitaxel (Fisher) was dissolved in DMSO to create a 5 mM stock solution, and diluted with half-strength Murashige-Skoog liquid media to a working concentration of 20 μM. Etiolated hypocotyls were transferred to petri dishes containing taxol in a dark room for 4h prior to imaging.

### Image processing and data fitting

Stacks of z-slices were merged using the Z-project function in ImageJ (http://rsbweb.nih.gov/ij/) (Schindelin et al., 2012). A cross-section of a z-stack was created using the Reslice function. Images were stitched together using the Stitching plug-in (Preibisch et al., 2009). To measure the MOR1-3xYPet microtubule distribution, the fluorescence intensity was measured using the RGB Line Profiler plug-in (http://rsb.info.nih.gov/ij/plugins/rgb-profiler.html). Specifically, 12 line-scans of 25 pixels in length were taken of microtubules and the fluorescence intensity of both YPet and mRFP were measured. The average intensity of each point and channel was measured. The background fluorescence was also measured and subtracted from the measured fluorescence of the line-scan. A similar method was used to measure MOR1-3xYPet length on microtubules.

To analyze the FRAP dataset, pre- and post-bleach intensity of the ROI, as well as the intensity of an unbleached region and a background region were measured in ImageJ (Schindelin et al., 2012). After subtracting the background signal, a correction was applied for incomplete bleaching such that the mean intensity of pre-bleached frames was set to 1 and the bleach frame intensity was set to 0. Using the ImageJ Curve Fitting tool (Schindelin et al., 2012), a least-squares fit (Pseudo-R^2^ cutoff = 0.75) was generated for the function F(t) = F_m_ * (1 - exp(-k_off_ * t)), where F_m_ is the mobile fraction and k_off_ is the off-rate of MOR1 on microtubules. Only the first 20 seconds immediately after bleaching were analyzed to minimize the effects of overall photobleaching and variable background signals.

### Microtubule growth rate analysis

Kymographs of individual MOR1 puncta were generated using the KymographClear macro toolset (Mangeol et al., 2016) in ImageJ. Only MOR1 on growing microtubules that could be tracked for a minimum of 10 sec was used for the analysis. Growth rates were calculated using the KymographDirect tool (Mangeol et al., 2016) or manually from kymographs.

The total number of tubulin additions per second was calculated based on the assumption that each α/β-tubulin dimer is ∼8 nm in length, and each microtubule has 13 protofilaments. For example, with MOR1-3xYPet in the *mor1-23* background at 21°C, microtubules grew at an average rate of 7.68 µm per minute, or 128 nm per second. Thus, ∼16 tubulin dimers were added to each protofilament or 208 tubulin dimers across 13 protofilaments every second. MOR1 residency time was calculated by 1/k_off_. Using the same example, each MOR1-3xYPet protein at 21°C stayed on the microtubule for an average of 3.6 seconds. Assuming 13 MOR1 proteins were active at the plus end, each MOR1 would have added, on average, 58 tubulin dimers.

### Protein structural modelling

TOG1 structures (amino acids 1-234) of MOR1 and mor1-1 (L174F) were predicted using Alphafold3 (Abramson et al., 2024) with *Arabidopsis* α-tubulin 4 (TUA4) and β- tubulin 4 (TUB4). Average hydrogen bond lengths within HEAT5 were calculated from five AlphaFold models in ChimeraX (Pettersen et al., 2021). Molecular dynamics of MOR1 and mor1-1 TOG1 without tubulin were simulated using the GROMACS package (ver. 2024.2) (Abraham et al., 2015, 2024) with the CHARMM36 force-field (Huang & MacKerell Jr, 2013). Proteins were solvated in a square box with TIP3P water molecules (Mark & Nilsson, 2001). The salt concentration was held constant at 0.15 M with NaCl in an explicit solvent model. Root mean square fluctuation (RMSF, in angstrom) at each residue of TOG1 was obtained with 2.5-nanosecond simulations at a constant temperature of 21°C and 30°C. All protein structures were visualized with ChimeraX (Pettersen et al., 2021).

### Statistical Analysis

Statistical analyses were performed with Microsoft Excel (Redmond, USA) or R (R Core Team, 2021) with the following packages: plyr (Wickham, 2011), FSA (Ogle et al., 2023), and rcompanion (Mangiafico, 2023).

## DATA AVAILABILITY

Research materials and experimental data will be available upon request.

## ACKNOWLEDGEMENTS

This research was funded by a Natural Sciences and Engineering Research Council of Canada (NSERC) Discovery grant (298264-09 and 2019-05432) to G.O.W., NSERC postgraduate scholarships to R.C.E. and L.S.H., a NSERC Undergraduate Summer Research Award to R.B., the Canada Research Chairs Program, and the Canada Foundation for Innovation. We thank the UBC Bioimaging Facility for confocal microscope access and assistance, Dr. Richard Cyr (Penn State University) for constructs and Dr. Ram Dixit (Washington University) for seed lines.

Molecular graphics and analyses performed with UCSF ChimeraX, developed by the Resource for Biocomputing, Visualization, and Informatics at the University of California, San Francisco, with support from National Institutes of Health R01-GM129325 and the Office of Cyber Infrastructure and Computational Biology, National Institute of Allergy and Infectious Diseases.

