## Supplemental Figures and Tables for "Microtubule plus-end dynamics are tightly coupled to the turnover of the MOR1 polymerase"

### Supplementary Figures

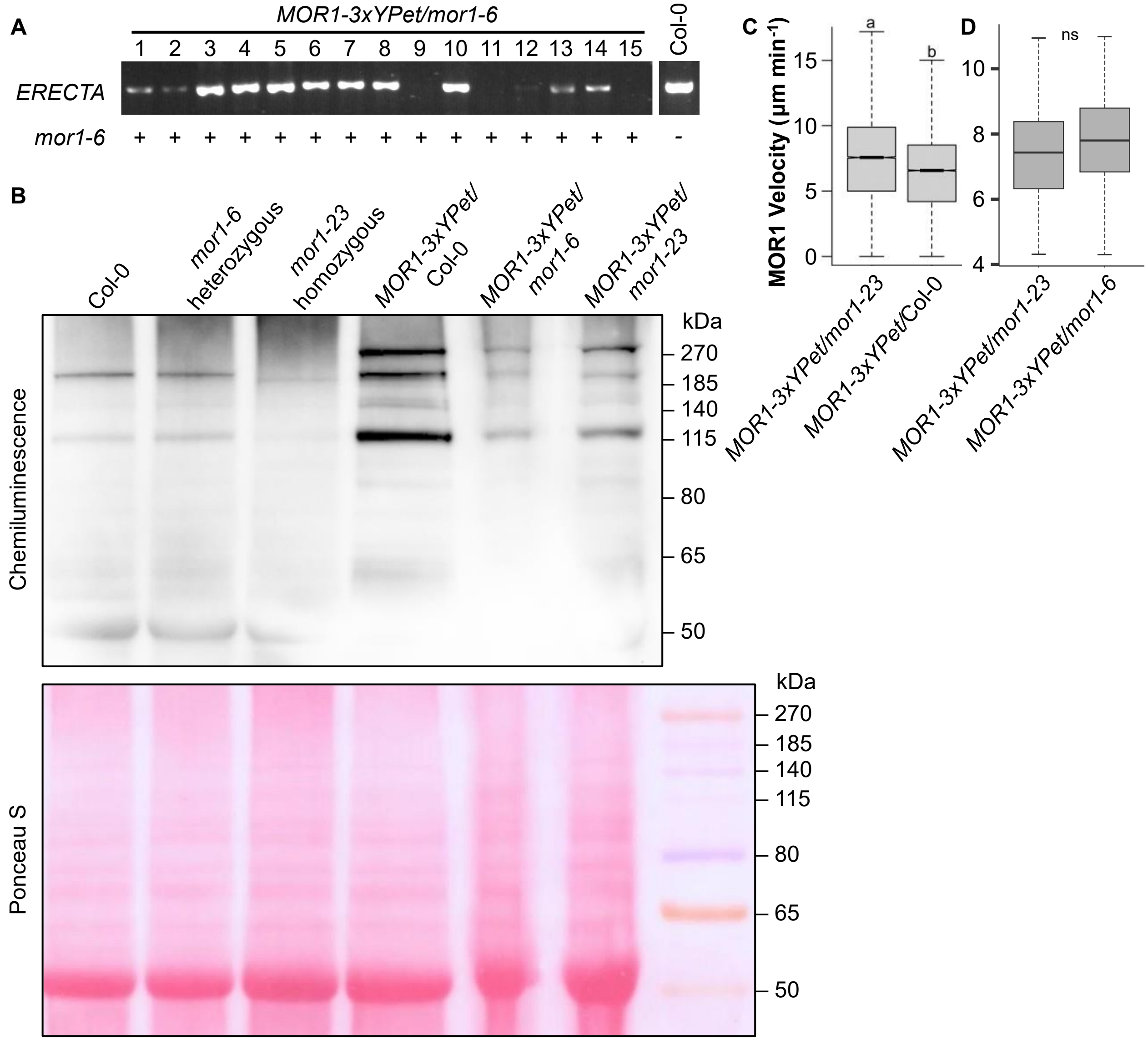


Supplementary Figure 1. Molecular analysis of Col-0, *mor1-23*, and *mor1-6* genetic backgrounds and the effect on MOR1 dynamics. **(A)** Genotyping results confirming the segregation of the wild-type *ERECTA* allele (presence of band) and the *er105* mutant allele (absence of band) in the *MOR1-3xYPet/mor1-6* line; as well as sequencing results confirming the presence (+) or absence (-) of the *mor1-6* point mutation. **(B)** Uncropped Western blot and Ponceau S-stained membrane confirming equal transfer of total proteins in each lane. **(C-D)** MOR1-3xYPet movements in elongated hypocotyl cells were imaged with TIRF at 21°C as a proxy for microtubule growth rate. **(C)** Increased *MOR1* expression in the Col-0 background led to slower MOR1 velocities (6.44 ± 0.03 µm min$^{-1}$) than in the *mor1-23* background (7.68 ± 0.03 µm min$^{-1}$). Mann-Whitney U-test $p < 0.001$, $n = 85$ particles for MOR1-3xYPet in the *mor1-23* background and $n = 80$ particles in the Col-0 background. **(D)** In a separate experiment, MOR1 velocities did not differ significantly in the *mor1-23* background (7.46 ± 0.19 µm min$^{-1}$) and in the *mor1-6* background (7.82 ± 0.18 µm min$^{-1}$). Mann-Whitney's U-test $p = 0.166$, $n = 64$ particles for both groups.

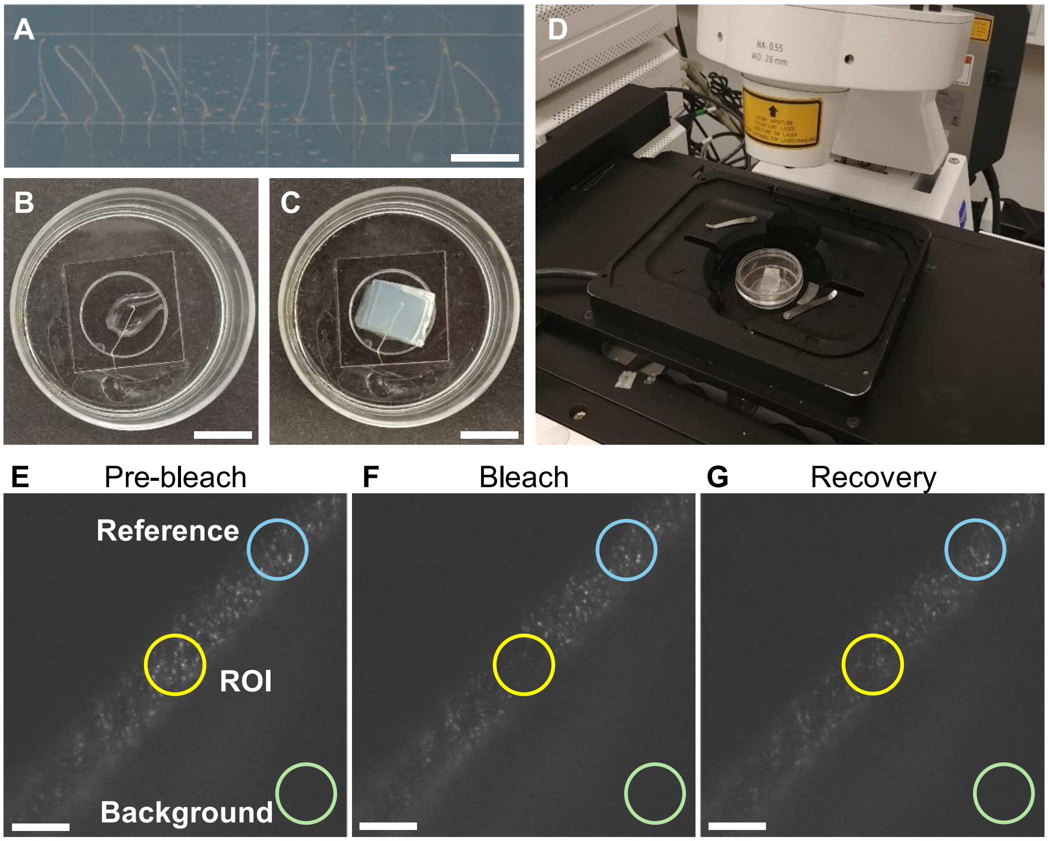


**H**

| ROI | REF | BACKGROUND | |
|---|---|---|---|
| 137.366 | 127.936 | 113.517 | Pre-bleach |
| 135.369 | 127.154 | 113.059 | |
| 135.833 | 126.418 | 112.866 | |
| 122.438 | 125.446 | 112.544 | Bleach |
| 122.667 | 124.474 | 111.985 | Recovery |
| 123.136 | 123.886 | 112.094 | |
| 123.456 | 123.98 | 111.897 | |
| 124.118 | 123.72 | 111.892 | |
| 123.841 | 123.875 | 112.013 | |
| 123.922 | 124.039 | 111.938 | |
| 123.976 | 123.562 | 111.926 | |
| 123.915 | 123.383 | 111.731 | |
| 124.141 | 123.101 | 111.8 | |
| 124.518 | 123.433 | 111.74 | |

**I**

| Time (s) | Intensity | |
|---|---|---|
| 0 | 0 | Bleach |
| 0.1 | 0.10086 | Recovery |
| 0.2 | 0.19332 | |
| 0.3 | 0.21639 | |
| 0.4 | 0.30421 | |
| 0.5 | 0.26257 | |
| 0.6 | 0.25482 | |
| 0.7 | 0.30641 | |
| 0.8 | 0.3179 | |
| 0.9 | 0.37078 | |
| 1 | 0.37165 | |

**J**

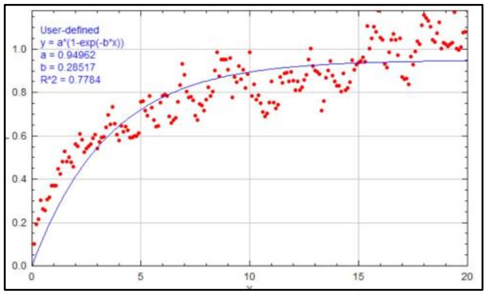


$$F(t) = F_m(1 - e^{-t \cdot k_{off}})$$

$$t_{\frac{1}{2}} = -\frac{\ln(0.5)}{k_{off}}$$

Supplementary Figure 2. TIRF imaging and FRAP analysis workflow.

**(A-D)** Mount seedlings for TIRF imaging.
**(A)** 4-day-old etiolated hypocotyls are grown on ½ MS media without sucrose. Scale bar = 1 cm.
**(B)** A circular dish with a cover slip on the bottom is used for imaging. The sample is immersed in liquid ½ MS (no sucrose) media, with the apical hook of the hypocotyl positioned in the centre of the cover slip. Scale bar = 1 cm.
**(C)** Three blocks of agar are placed on top of the hypocotyl to gently press the specimen closer to the cover slip. Scale bar = 1 cm.
**(D)** The mounted specimen is placed on the stage of the Zeiss TIRF microscope.

**(E-G)** Define regions of fluorescence quantification for FRAP analysis. In each image, the fluorescence is recorded from the region of interest (ROI, yellow circle), an area within the cell beyond the ROI to account for gradual photobleaching over time (Reference, blue circle), and an area outside of the cell to quantify the background fluorescence (Background, green circle). Scale bars = 10 μm
**(E)** Acquire time-lapse videos. Make sure to acquire at least three frames prior to photobleaching.
**(F)** A pulse of laser is used to photobleach fluorophores in the ROI.
**(G)** Images are captured following the photobleach to measure how quickly recovery of fluorescence
takes place.

**(H-J)** FRAP modelling.
**(H)** Use FIJI to quantify fluorescence intensities within each of the three regions (ROI, reference, background) over time. Compile intensities into three columns in a spreadsheet.
**(I)** Use software (such as R) to calculate normalized fluorescence recovery over time. The normalized intensity of the bleached frame is set to zero.
**(J)** Use software (such as FIJI) to generate a best-fit curve through the data points. Values of mobile fraction ($F_M$) and $k_{off}$ are obtained from curve fitting.

### Supplementary Tables

Supplementary Table S1. Primers used in this study.

| Primer Name | Primer Sequence (5' to 3') | Function |
|---|---|---|
| *At2g35630*F | GTCTTCAAGCCCGAATGGAGAGGCTCAAAGGTGG ATCACTGGAACATATG-GGAGGTGGAGGTGGAGCT | Recombineering |
| *At2g35630*R | GAATTTGCTTATGTAACTTCAGACTCCAAAGCTTG GGGTCTCTAGTTCTA-GTGATGGTGATGGTGATG-GGCCCCAGCGGCCGCAGCAGCACC | Recombineering |
| *At2g35630*TestF | GGACGCTAGATGCAATCAGAG | Recombineering |
| *At2g35630*TestR | GAATCTGGCACACTCACCATC | Recombineering |
| *At2g35630*LBdel | ATGATTTTGTGTGGTACCACAAGATCAACCGCTTC AAAAACATAGAACAA-TAGGAACTTCCCCCTCTTGG | Recombineering |
| replaRB-amp | TATATTGCTCTAATAAATTTTTGGCGCGCCGGCCA ATTAGGCCCGGGCGG-TTCAAATATGTATCCGCTCATG | Recombineering |
| *At2g35630*RBDel | GGAGAGAATATTTCTCAATGTCTCCTCTCCCAATC TTCCATCGTCTTCCT-TTACCAATGCTTAATCAGTG | Recombineering |
| replaLB-tet | TTAGTTGACTGTCAGCTGTCCTTGCTCCAGGATGC TGTTTTTGACAACGG-TAATTCCTAATTTTTGTTGACAC | Recombineering |
| *At2g35630*TestDelLB | AAGAAAATATCCAGTTGACTCC | Recombineering |
| *At2g35630*TestDelRB | TACAAAAGCAGCTCGACATGC | Recombineering |
| RBtest | AAATATAGCGCGCAAACTAGG | Recombineering |
| LBtest | CTGAATTCTTAATTAACCCACG | Recombineering |
| *mor1-1*GalKF | TTAGTGAATTTGGATCAAAAGTTATTCCACCTAAAA GGATTTTAAAGATG-T-GGAGGTGGAGGTGGAGCT | Recombineering |
| *mor1-1*GalKR | TTTGGCAGATGCACGGACATTCTGATCTTGATGGT CGAAAAGTTCAGGAA-GGCCCCAGCGGCCGCAGCAGCACC | Recombineering |
| *mor1-1*repF | TTGCAATAACTCTACTTGATATGC | Recombineering |
| *mor1-1*repR | TTTGGCAGATGCACGGACATTCTGATCTTGATGGT CGAAAAGTTCAGGAA-A-CATCTTTAAAATCCTTTTAGG | Recombineering |
| *mor1-1*testF | TGGGTAGAATTAGAGGCCGTCG | Recombineering |

| **Primer Name** | **Primer Sequence (5' to 3')** | **Function** |
|---|---|---|
| *mor1-1*testR | CTCAACATAAATCTTACCATCG | Recombineering |
| SALK_032056 LP | TTCAACAGCCAACAATCCTTC | *mor1-23* genotyping |
| SALK_032056 RP | GAATCTGGCACACTCACCATC | *mor1-23* genotyping |
| LBb1.3 | ATTTTGCCGATTTCGGAAC | *mor1-23* genotyping |
| *er105* fwd | TCTGTCGTCTGAGCACTCAAT | *er105* genotyping |
| *er105* rev | AGTTGTTTCCTCGCAACCCA | *er105* genotyping |
| *mor1-6* seq fwd | GGATTGCCTTGATTTGTCTAG | *mor1-6* sequencing |
| *mor1-6* seq rev | CTGAGGTAGATAACTCAAGTA | *mor1-6* sequencing |
| *MUSE3* fwd | GTGGGAGCAGAGACCAACTC | qRT-PCR |
| *MUSE3* rev | TGGCAACCCTCTCAACCATC | qRT-PCR |
| *MOR1* fwd | TTTGCGATGCTATTGCGCTC | qRT-PCR/dPCR |
| *MOR1* rev | CTCCATCGTATCCAGAAAAACATCG | qRT-PCR/dPCR |
| *mor1-6* cDNA fwd | CTGCGGCTCCAAAGAGAGTT | RT-PCR |
| *mor1-6* cDNA rev | GCACCAAGCCACAAGTCAAG | RT-PCR |
| *mor1-23* cDNA fwd | TCCCCCTTCGTCCTTAGCTC | RT-PCR |
| *mor1-23* cDNA rev | TCTGGCACACTCACCATCAG | RT-PCR |
| *CLASP* DT1-BsF | ATATATGGTCTCGATTGTGCGTATTTCAGCTATGC GGGTT | CRISPR/Cas9 |
| *CLASP* DT1-F0 | TGTGCGTATTTCAGCTATGCGGGTTTTAGAGCTAG AAATAGC | CRISPR/Cas9 |
| *CLASP* DT2-R0 | AACCCGCATAGCTGAAATACGCACAATCTCTTAGT CGACTCTAC | CRISPR/Cas9 |
| *CLASP* DT2-BsR | ATTATTGGTCTCGAAACCTGGAAATTGGGAACATG GCC | CRISPR/Cas9 |

**Supplementary Movies**

Supplementary Movie S1. MOR1 is a plus-end tracking protein.
Time-lapse of spinning-disc confocal images of MOR1-3xYPet (in red) plus-end tracking on growing microtubules (in cyan). MOR1-3xYPet particles are labelled with yellow circles. Images were acquired every 5 seconds. Scale bar represents 20 μm. Movie is 5 frames per second.

Supplementary Movie S2. MOR1 localizes to catastrophe and rescue events of microtubule plus ends.
Time-lapse of spinning-disc confocal images of MOR1-3xYPet (in red) localizing to a microtubule plus end under catastrophe and rescue (in cyan). Growing plus ends and shrinking plus ends are labelled with arrowheads and arrows, respectively. Catastrophe of microtubule occurs at 75 seconds and rescue occurs at 90 seconds. Images were acquired every 5 seconds. Scale bar represents 5 μm. Movie is 5 frames per second.

Supplementary Movie S3. MOR1 is not found on microtubule minus ends.
Time-lapse of spinning-disc confocal images of a treadmilling microtubule (in cyan). MOR1-3xYPet (in red) is not found on minus ends. Growing plus ends and shrinking minus ends are labelled with arrowheads and circles, respectively. Images were acquired every 5 seconds. Scale bar represents 5 μm. Movie is 5 frames per second.

Supplementary Movie S4. MOR1 localizes to newly created plus ends from microtubule-severing events at cross-over sites. Time-lapse of spinning-disc confocal images of a microtubule undergoing severing following a crossing over event. MOR1-3xYPet localized to a growing microtubule plus end (labelled by the arrowhead) is followed by a severing event at 30 seconds (labelled by an arrow) following a crossing-over event. This is followed by MOR1 localization at the depolymerizing plus end (labelled by a circle). Images were acquired every 5 seconds. Scale bar represents 5 μm. Movie is 5 frames per second.

Supplementary Movie S5. MOR1 localizes to mitotic microtubule arrays. MOR1 localizes to microtubules from the pre-prophase stage to late anaphase stage. Time-lapse spinning-disc confocal micrographs of MOR1-3xYPet (in cyan) and mRFP-TUB6 (in red) being expressed in dividing cells in the root tip. MOR1-3xYPet localizes to microtubule structures in all stages of mitosis. Each individual image shows two cells at different stages of mitosis and is either labelled with a white or yellow bracket. The white bracket labels a cell starting from the pre-prophase to anaphase stage. Note that MOR1-3xYPet localization occurs near the nuclear envelope during its breakdown followed by localization to the mitotic spindle. The yellow bracket labels a cell that has progressed further in mitosis from metaphase to late anaphase. Time progresses from left to right. Images were acquired every 60 seconds. Scale bar represents 10 μm. Movie is 5 frames per second.

Supplementary Movie S6. Representative MOR1 FRAP movies under various experimental conditions. **(A)** MOR1-3xYPet in the Col-0 background, 21°C. **(B)** MOR1-3xYPet in the *mor1-23* background, 21°C. **(C)** MOR1-3xYPet in the *mor1-23* background, 30°C. **(D)** *mor1-1-YPet* in the *mor1-23* background, 21°C. **(E)** *mor1-1-YPet* in the *mor1-23* background, 30°C. **(F)** MOR1-3xYPet in the *mor1-23* background, 21°C with taxol. **(G)** MOR1-3xYPet in the *mor1-23* and the CRISPR *clasp* backgrounds, 21°C. Movies were collected from 3- to 4-day-old etiolated hypocotyls at 100 ms/frame for 20 seconds. Scale bar represents 5 μm.
